# Organelle Development and Inheritance are Driven by Independent Nuclear and Organellar Mechanisms in Malaria Parasites

**DOI:** 10.1101/2025.03.06.641809

**Authors:** Shahar Michal, Qasem Alia, Freedman Eshkar, Maron Yariv, Tissawak Amanda, Florentin Anat

**Affiliations:** The Kuvin Center for the Study of Infectious and Tropical Diseases & Department of Microbiology and Molecular Genetics, Faculty of Medicine, The Hebrew University of Jerusalem, Israel; Core Research Facility, Faculty of Medicine, The Hebrew University of Jerusalem, Israel

**Keywords:** malaria, *Plasmodium falciparum*, parasitology, apicoplast, cell cycle, DNA replication, ploidy, dynamic imaging

## Abstract

The apicoplast organelle of *Plasmodium falciparum* is essential for parasite’s replication, however the details of its biogenesis, inheritance and regulation throughout the cell cycle are unknown. Here, we report the development of a dynamic imaging platform coupled with an analytical pipeline that enables us to follow and measure subcellular structures throughout the 48-hour cell cycle of live parasites. We reveal a predetermined sequence of four discrete morphological steps in organelle development, which are tightly correlated with nuclear replication. We show that one of these steps, which we term the Crown morphology, is required for nucleus-apicoplast attachment. During Crown, apicoplast is stretched over multiple nuclei, fastened by centriolar tubulin. A complementary molecular approach was used to discover the basic ploidy of apicoplast and mitochondrial genomes, their replication rates and association with nuclear DNA replication. We inhibited nuclear DNA replication and found that it completely blocks apicoplast biogenesis in its most initial stages, demonstrating dependency on S-phase initiation. Conversely, specific inhibition of apicoplast genome replication resulted in an almost-undisturbed organelle development and division. However, it affected the Crown step, preventing association to tubulin-nuclear structures, leading to failure in accurate organelle sorting into daughter cells. Collectively, these experiments reveal a central cellular pathway linking apicoplast development to the parasite’s cell cycle, and a second independent organellar mechanism responsible for segregation into daughter cells.

## Introduction

The deadliest malaria parasite, *Plasmodium falciparum*, possesses a unique subcellular organelle called apicoplast^1,2^. This four-membrane bound compartment originated through an ancient secondary endosymbiotic event, in which a protist engulfed a red alga together with its chloroplast^3^. Although all photosynthetic abilities were lost during evolution, the apicoplast retained essential metabolic pathways that make it an indispensable cellular entity throughout the parasite’s complex life cycle^4^. Key Functions include the biosynthesis of isoprenoids, Heme, iron-sulfur clusters, fatty acids, and coenzyme A^5^. These metabolic pathways are carried out by enzymes with a prokaryotic origin and plants-like characteristics. Prokaryotic-like functions also underlie most other known apicoplast housekeeping functions, such as biogenesis, replication and proteostasis^3^. Consequently, several clinical antimalarial drugs target the apicoplast, and many more apicoplast-targets could be identified with a better understanding of apicoplast biology^6,7^.

Almost all apicoplast resident proteins, enzymes included, are encoded by the cell nucleus and are transported to the organelle through the secretory pathway. However, similar to other endosymbionts such as mitochondria and chloroplasts, the apicoplast also carries its own genome, in the form of a small circular chromosome. Besides sequence and computational predictions, not much is known of the apicoplast DNA^8^. Basic facts such as copy number, replication mechanisms and timing or are limited or missing. This genome is predicted to encode 68 genes, the vast majority of those are thought to be related to organellar protein synthesis^8^. Consequently, clinical antimalarials such as doxycycline are effective due to their inhibitory effect on protein translation in the apicoplast^9,10^. Any other regulatory functions of the apicoplast genome remain to be discovered.

The asexual replication of *P. falciparum* inside the red blood cell (RBC) begins with invasion by a single parasite into the host RBC and culminates 48 hours later in the egress of roughly 30 new daughter parasites^11^. Unlike human and plant cells, the parasite has a single mitochondrion and a single apicoplast in each cell and during this unique cell cycle, they undergo dramatic morphological changes^12^. Early in the cycle, the apicoplast is found in a small globular shape and remains this way throughout most of the parasite’s intraerythrocytic development. When the cell nucleus begins to replicate and divide, the apicoplast starts to elongate, branch, and divide^13^. These morphological changes are typically documented using images of fixed and live cells, providing snapshots of different stages in organelle biogenesis^14–16^. Two recent studies shed light on common features in the divisions of mitochondrion and apicoplast. Verhoef and colleagues used ultra-resolution imaging to identify close association with the mitochondrion^17^, whereas Morano *et al* used a combination of genetics and live imaging to show a common fission mechanism for both organelles^18^. Yet, a complete sequence of events including the entirety of organelle morphologies, their physical properties, transitions and kinetics is currently missing. Moreover, most of the molecular mechanisms governing these processes are still unknown. Importantly, during its curious cell cycle *Plasmodium* replicates its nucleus multiple times, giving rise to dozens of daughter cells in a process called schizogony. These nuclear replications are asynchronous and result in a variable number of progenies^19^. It is therefore completely unclear how the apicoplast senses the final number of daughter cells to guarantee accurate division and sorting.

Here, we developed a platform for dynamic imaging, and discovered new morphologies and kinetics of nuclear replication and organelle biogenesis in high resolution. We further established an analytical pipeline to quantify changes in organelles’ discrete developmental steps. Using molecular tools, we determined the copy number of organelles genome and their replication rates. We used a pharmaceutical approach to reveal that apicoplast biogenesis completely depends on nuclear replication. Conversely, apicoplast growth proceeded normally regardless of organelle genome replication, with the exception of a single developmental step. Finally, we provide initial evidence for a putative role of tubulin in accurate organelle segregation into daughter cells and its link to organelle genome inheritance. Collectively, these experiments reveal a central cellular mechanism linking apicoplast development to the parasite’s cell cycle, and a second independent organellar mechanism responsible for segregation into daughter cells.

## Results

### Establishing an experimental platform for dynamic imaging of live *Plasmodium* parasites

To follow organelle development and cell cycle progression, we sought a viable, fluorescent-tagging method for co-live-tracking of the nucleus and the apicoplast. Unlike many eukaryotic cells, nuclear division in the parasite occurs via closed mitosis inside an intact nuclear envelope^19,20^. These multiple nonsynchronous replications within a miniscule space set hurdles for following individual nuclei during schizogony. We therefore chose an H2B orthologue^21^ (PF3D7_1105100) with the prospect of overcoming signal diffusion by close association with nuclear DNA. For apicoplast tracking, we used the first 60 amino acids of acyl carrier protein, a known and well-studied apicoplast residing protein^22^. This N-terminus comprises a transit peptide (TP) which we and others have previously used to guide fluorescent proteins into the apicoplast matrix^22–24^.

The construct H2B-mRuby-2TA-TP-mNeon was inserted into the parasite genome using the attB/attP transposon-mediated integration system^25^ (Fig. 1A and Fig.S1). Live microscopy was used to verify the expression and localization of the green and red signals and the overlap of the H2B-red signal with a DNA staining throughout the parasite’s cell cycle (Fig. 1B). As recently published^17,18^, we observed the beginning of apicoplast elongation at the onset of schizogony, and its division into individual organelles at the end of it (Fig. 1B). Next, we set out to design a viable protocol that could provide a detailed description of subcellular events throughout the parasite asexual growth, with a focus on nuclear divisions and apicoplast biogenesis. To do that, we optimized a dynamic imaging platform that supports long term imaging (>24h), with short time intervals (15 min) and high-resolution three-dimensional image acquisition (Z<0.5 μm) (Fig. 1C). The parasite’s acute sensitivity to oxidative stress^26^, made phototoxicity the most critical challenge to overcome. Viability was determined by the ability of imaged parasites to egress and reinvade. Consequently, system optimization primarily mitigated those issues at the sample preparation and microscopy steps along with automatization of image analysis (Fig. 1C and Methods).

**Figure 1.**
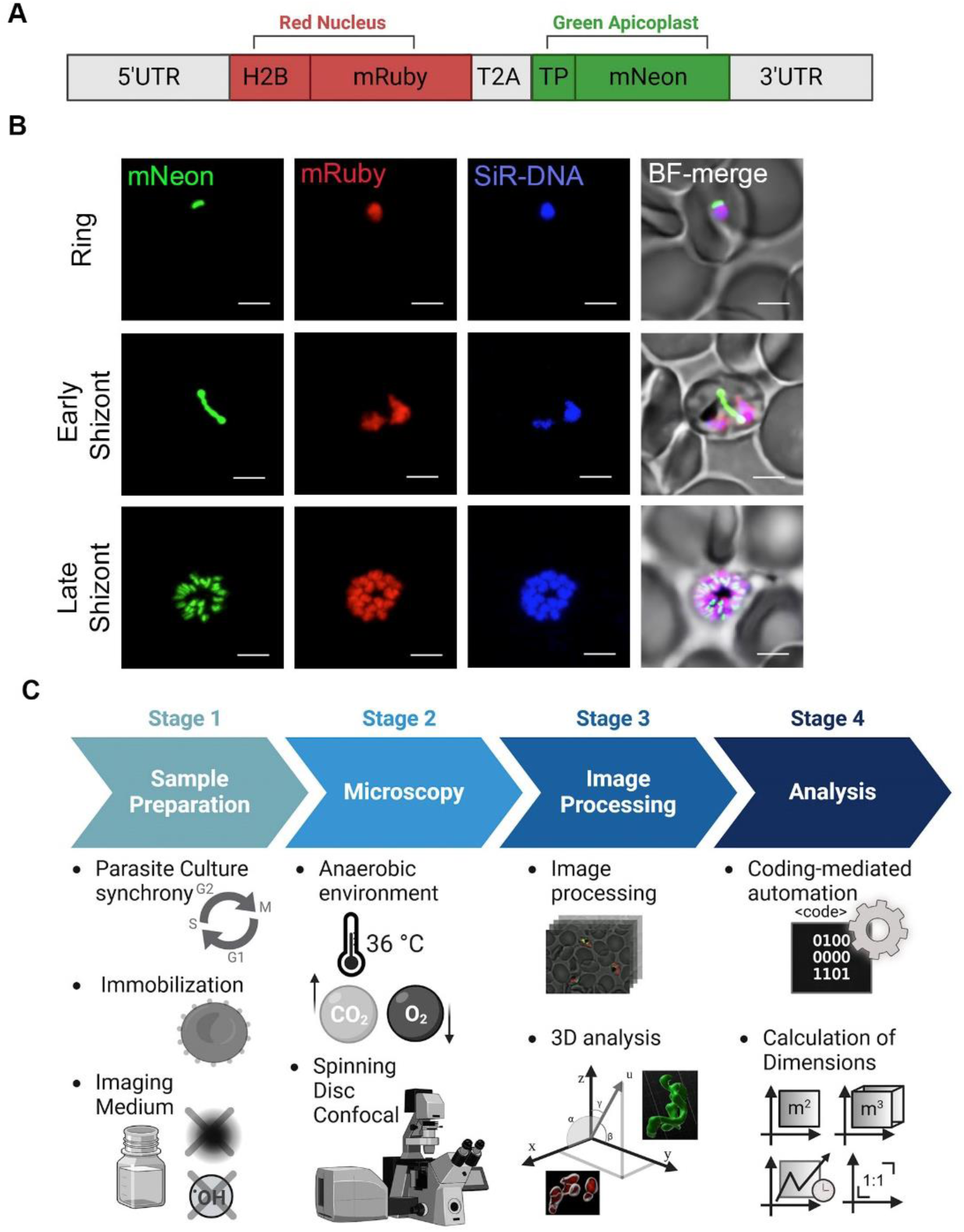
Establishing a dynamic imaging platform to follow subcellular events throughout parasite’s 48h life cycle. **A.** Schematic representation of the genetic reporter. mRuby was fused to histone H2B for nuclear labeling, while mNeon was linked to an apicoplast-specific transit peptide (TP) for organelle targeting. The T2A sequence facilitates ribosomal skipping during translation, enabling the expression of two separate proteins driven by a single promoter. **B.** Representative fluorescence live microscopy images of the H2B-mRuby/TP-mNeon transgenic parasite line at different stages of its life cycle. Images were captured as Z-stacks (total 21 slices of 0.5 μm each) at 100x and are displayed as maximum intensity projection. Apicoplast is green, and nucleus is distinctly labeled in red. DNA staining with SiR-DNA (blue) confirms nuclear localization of mRuby signal. Scale bar is 2.5 μm. C. Workflow for long-term live imaging and biogenesis analysis, highlighting sample preparation and microscopy optimization, followed by automated image processing and analysis.

### Four distinct apicoplast developmental stages are identified during Schizogony

In an attempt for dissection of sequential organellar events, we imaged parasites throughout the entirety of their 48-hour cell cycle. As expected^16^, during the first half of the cell cycle the parasites are metabolically active, but no morphological changes can be microscopically observed in their single haploid nucleus or their single globular apicoplast (Fig.S2A). We then focused our attempts on the second half of their cell cycle, imaging parasites throughout the last 24 hours. We verified the viability of imaged parasites by their ability to egress and reinvade new RBCs, determining egress as the 48h time-point (Fig. 2A). Consequently, all sequential events were counted-back from egress and were put on the cell-cycle timeline. Thus, we observed that the first change in apicoplast shape begins roughly 16 hours before egress, at 32 hours post invasion (Fig. 2A). At this time point, the so-far globular structure, roughly 0.5 μm in diameter, begins to elongate. In the following hours the organelle undergoes extreme morphological changes, culminating in its division and sorting into individual daughter cells. We observed four distinct steps that the apicoplast undertakes in the last third of the cell cycle: (1) Elongation; (2) Branching; (3) Crown; (4) Division (Fig. 2A and Sup. Movie1, see next section for more details). While a 2D visualization of maximal projection provided essential details of these structures, 3D analysis further enhanced characterization and contributed to downstream investigation (Fig. 2A. bottom panel). The complete 3D structures of the organelles were constructed from twenty Z-steps of 0.5 μM each. Physical parameters of these subcellular structures were extracted and analyzed via different methodologies, including 3D Suite plugin for Fiji^27,28^ and IMARIS package (Fig. 2B and Sup. Movie2) (see methods for details). Importantly, these images were comparable with what could be observed with expansion microscopy (Fig.S2B). Furthermore, these images resemble 3D rendering done by volume electron microscopy^12,17^, while avoiding potential fixation artifacts and gaining kinetic data.

**Figure 2.**
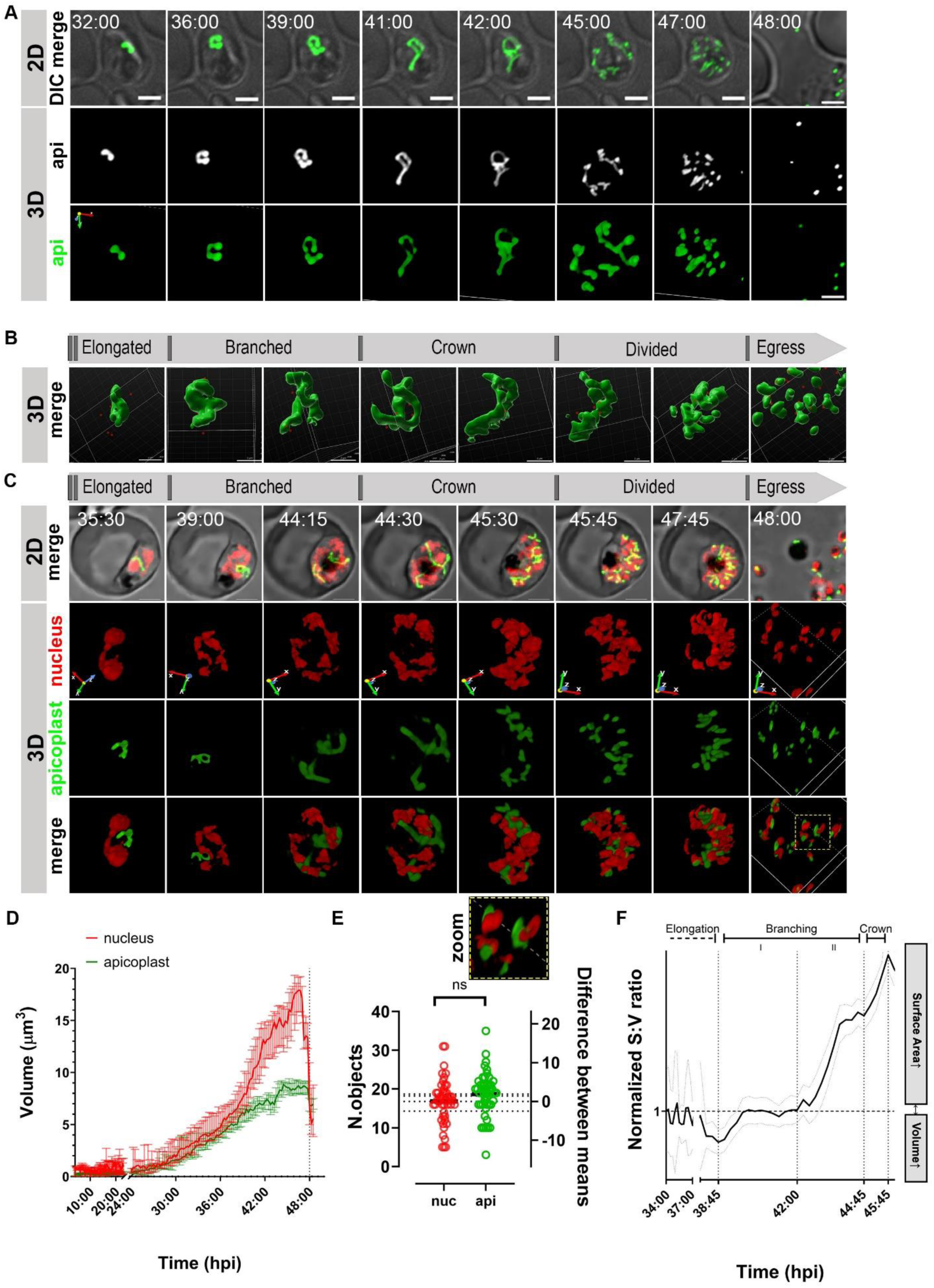
Dynamic imaging reveals morphological details, kinetics, physical measurements and a tight synchrony in the co-development of nuclei and apicoplast. **A.** Representative images from sequential capturing of apicoplast biogenesis of *P. falciparum.* Images were captured as Z-stacks (total 21 slices of 0.5 μm each) at 100x for 20h with 30 min intervals using spinning disc confocal microscope. The images are displayed as maximum intensity projections (2D) or volume reconstructed for 3D visualization using NIS image analysis software. Scale bar is 2.5 μm. **B.** 3D reconstructed images of H2B-mRuby/TP-mNeon parasites using Imaris image analysis software show detailed apicoplast morphologies (green) along with nuclei location (red dots) during life cycle. Scale bars, 2.5 μm. **C.** Representative long-term fluorescence live-cell microscopy images show detailed apicoplast (green) and nucleus (red) morphologies and conformations during different stages of parasite’s life cycle in 2D (as maximum intensity projections of Z-stack images) and 3D (volume visualizations). Microscopy Technique: Spinning disk confocal microscopy. Images were captured at 100x as Z-stacks (total 21 slices of 0.5 μm each) with 15 min time intervals for 20 h. Scale bars, 2.5 μm. **D.** Quantification of median organelle volume over time was obtained as described in Materials and Methods for 151 parasites (n>20 for each time point), from 4 independent experiments. Error bars are 95% CI. **E.** Final number of organelles per parasite was obtained as described in Materials and Methods for n>40 parasites from 4 independent experiments. The difference between nuclei and apicoplast mean number of objects (17+/-5.8 (nuc) and 19 +/- 5.6 (api), (api - nuc) =2 +/- 3.8) is not significant. Unpaired Mann Whitney test, P =0.06 or Unpaired t test with Welch’s correction, P=0.1, ns. **F.** Quantification of median organelle surface area: volume ratio vs. time was obtained as described in Materials and Methods for n>20 parasites for each time point, from 4 independent experiments. Surface area to Volume ratio vs. time is a mathematical representation of the extensive branching and shape-shifting of the apicoplast, particularly expanding at the last 6 h of the cell cycle.

We then proceeded to dual-labeling 4D live microscopy and analyzed the co-development of the apicoplast (green) and the nucleus (red). We observed that the first nuclear division (one to two nuclei) coincides with the beginning of Elongation (step 1) at roughly 32 hours post invasion (Fig. 2C, Fig.S2C and Sup. Movie3). Apicoplast Elongation (step 1), characterized by five hours of linear growth, coincides with the first two to three rounds of nuclear replications. Detection of individual nuclei was possible during Elongation as well as immediately before and post-egress (Fig. 2C). Despite using the histone marker and the high image quality, we could not separate individual nuclei in the last ten hours before egress, during steps (2), (3), (4) of apicoplast biogenesis (Fig. 2C and Sup. Movie4-6). This points to tight packing and close connections between nuclei during schizogony as previously suggested^19^. Importantly, close proximity between nuclei and expanding apicoplast is observed during the late stages of apicoplast biogenesis. During the six hours of organellar bifurcating events in Branching (step 2), apicoplast is only partially associated with nuclei. However, a close association between the yet-undivided apicoplast and all divided nuclei was observed during the short Crown phase (step 3) (Fig. 2C). The Crown morphology is typically characterized by a closed circle with minute budding structures formed just prior to organelle division (Fig. 2A). Collectively, apicoplast biogenesis could be defined by four distinct developmental steps. These defined morphological structures physically entwine the replicating nuclei during schizogony in a highly coordinated matter.

### Measuring kinetics and physical properties of apicoplast-nuclear co-development

To quantify these events, we used the raw 3D data to calculate total volume of both organelles throughout the entire cycle. Combining multiple experiments, we measured the volumes of a total of 151 individual cells (Sup. Movie7), and observed a tight synchrony between subcellular events (Fig 2D). These averaged volume measurements confirm that no growth occurs in the first half of the cycle. Roughly 32 hours post invasion, both organelles begin to grow, the nuclei through replication and the apicoplast through Elongation. Strikingly, in the next ten hours during Elongation (1) and Branching (2), the two grow at the same pace, almost to the same dimensions, demonstrating tight temporal and spatial synchronizations. In the last eight hours of the cycle, the growth rate of nuclei volume exceeds that of the apicoplast. The nuclei collectively increased their total volume by roughly 20 folds, whereas the apicoplast plateaued out at 9-fold of its initial volume (Fig. 2D). The sharp drop at the 48-hours timepoint represents egress and the point in which individual nuclei can be deciphered. At this time point, when we counted individual organelles, we observed an average of 20 nuclei and 20 apicoplast organelles (‘objects’) per egressed cell. These represent twenty daughter cells egressing from a ‘mother-cell’, each equipped with a single nucleus and a single apicoplast, ready for reinvasion (Fig. 2E and magnified view). The rapid 9-fold growth in apicoplast volume during Elongation and Branching, in theory, could have resulted in a drop of surface area: volume (SA:V) ratio. This would have posed a serious challenge for an organelle required for metabolic exchange with the engulfing cell. We therefore measured and quantified this ratio throughout the last 14 hours before egress, and observed that not only does it stay constant, but also increases at specific steps (Fig. 2F). These changes in SA:V ratio directly result from the shapeshifting behavior of the apicoplast and therefore could be used as a proxy for defining the four developmental steps (Fig. 2F): (1) Elongation – five hours of linear growth, with a constant SA:V ratio and a final minor drop. (2) Branching – six hours of splitting and diverging structures, with a phase I of a constant SA:V ratio, and a phase II of a rise in SA:V ratio. And (3) the Crown: a short one-hour step, characterized by a sharp increase in SA:V ratio (Fig. 2F). The rounded, typically closed Crown morphology formed prior to organelle division, is easily missed or overlooked in fixed samples (Fig. 2A). As will be demonstrated below, the Crown morphology appears to be a key step towards accurate sorting of daughter organelles after division.

### Tubulin links apicoplast to nuclei during the Crown step

We next looked into the mechanism of accurate apicoplast division, and the sorting of individual apicoplast organelles into merozoite daughter cells. Results presented above suggest that such mechanism may go into action during the Crown step, as it seems to involve association between the yet-undivided apicoplast and the individual nuclei. We further hypothesized that this mechanism involves a cytoskeletal element, and decided to use a tubulin live-stain. This enables us to follow the dynamics of the *Plasmodial* centrosomes, often called centriolar plaques (CPs)^29^. We optimized a protocol for viable tubulin-tracking and followed tubulin association with the nuclei and apicoplast in the last five hours before egress (Fig. 3A and Sup. Movie 8). As previously documented, mitotic-spindle microtubules are associated with the nuclei during schizogony and, together with the CP machinery, provide the pulling forces for mitosis^14,29^. Using dynamic imaging, we observed that during Elongation and Branching stages, a single tubulin structure was associated with each individual nucleus, but not with the intricate apicoplast structure (Fig. 3B and Sup. Movie 8). However, in the very last stages before division, the process of Crown formation begins and the apicoplast gradually extends over multiple nuclei. As apicoplast stretches over nuclei, a structured and interconnected arrangement is formed. The apicoplast conforms to the contours of nuclei distribution in space, defining in each cell its unique Crown morphology. The attachment is secured by tubulin, which acts as a fastening component, strategically linking the yet-undivided apicoplast to multiple nuclei. As a result of a Crown-like formation, each tubulin structure becomes associated with the apicoplast on one end and a single nucleus on the other (Fig. 3A, 3B and Sup. Movie 9). Interestingly, the nuclei and tubulin do not change much their spatial arraignment since the late Branching step, reaching their final localization six hours before egress (Fig. S3). The apicoplast, however, becomes associated with tubulin-nuclei structures only three hours later, during the Crown stage. Thus, apicoplast Crown morphology is determined by the prior arrangement and numbers of nuclei-tubulin pairs (Fig. S3). This symmetric arrangement formed by the three subcellular compartments is followed by an accurate division with equally segregated daughter cells (Fig. 3 and Sup. Movie 8).

**Figure 3:**
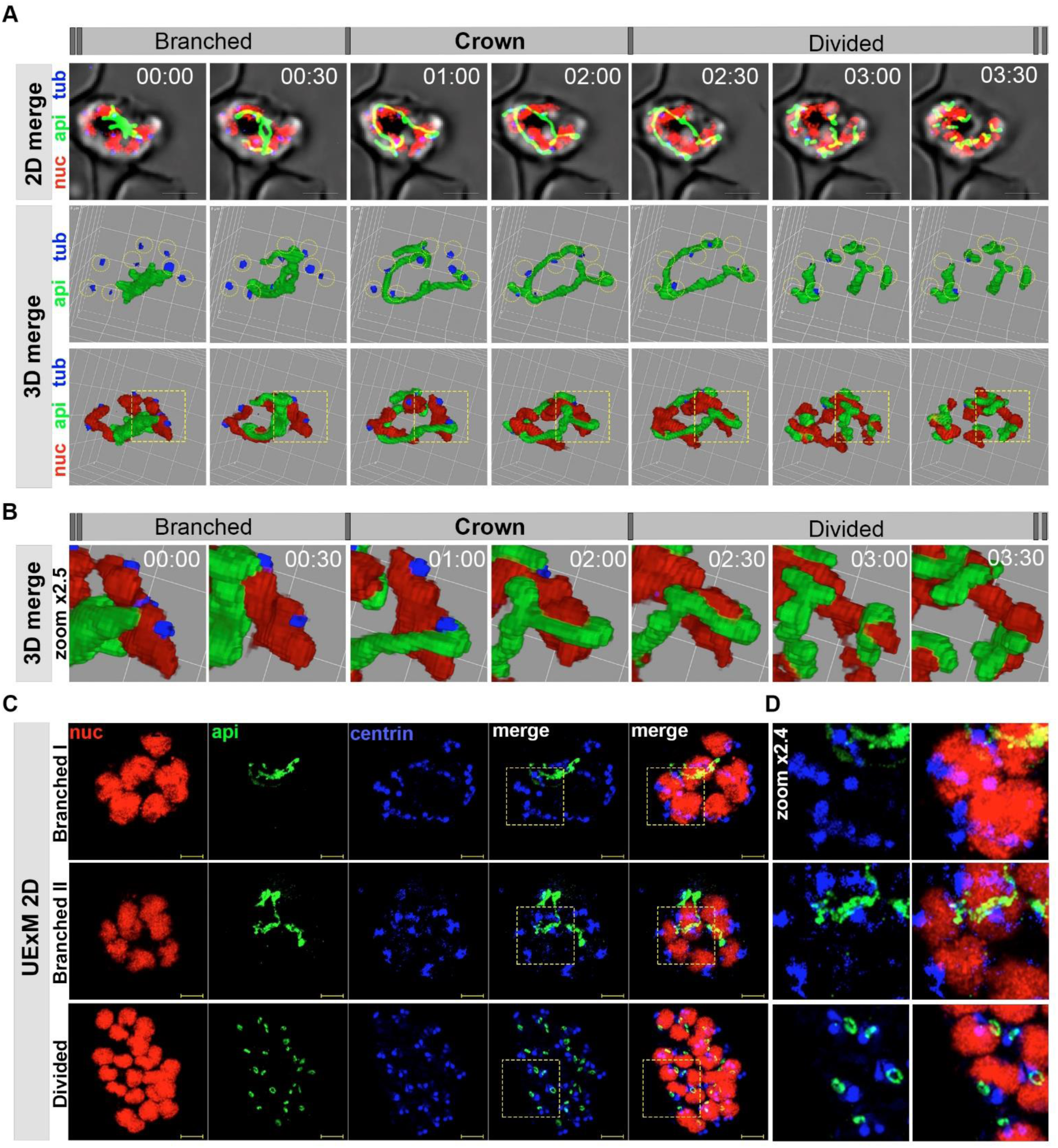
Tubulin links apicoplast to nuclei during the Crown step. **A.** Representative fluorescence live long-term microscopy images of whole-cell parasite reporter line H2B-mRuby/TP-mNeon stained with Tubulin Tracker (blue), show detailed apicoplast (green) and nucleus (red) morphologies and conformations during final stages of parasite’s life cycle in 2D (maximum intensity projection) and 3D (volume visualization) using NIS image analysis software. In untreated parasites, dots of tubulin structures observed next to each nucleus from early replication stages, whereas the apicoplast is not co-localize with tubulin structures. Later at the crown stage, the apicoplast is aligned over nuclei and the tubulin structures. Scale bars, 2.5 μm. 3D grid 2×2 μm. **B.** Images of individual CP pairs provide a magnified view of the cell shown in (A), illustrating the timeline of tubulin assembly—first with the nuclei and later with both the nuclei and apicoplast. Since tubulin is positioned between the nuclei and apicoplast, it becomes obscured when assembled in this region. **C.** Representative images of expanded parasites from different cell-cycle stages demonstrate centrin association with nucleus and later with both the nuclei and apicoplast. Apicoplast (green), nucleus (red) and centrin (blue) were stained as described in Ultra-Structure Expansion Microscopy section in Materials and Methods. Images were captured as Z-stacks (10-21 slices of 0.13 μm each) at 63x using Airyscan microscopy. The images are displayed as maximum intensity projections. Scale bar is 5 μm. D. Magnified views of the marked areas in (C) illustrate the progressive association of centrin with both organelles.

To further investigate the nature of these apicoplast-nuclei contact sites, we looked directly at the CPs at the nuclear envelopes. To do that, we used an anti-centrin antibody, an established marker for the highly divergent *Plasmodium* centrosome^14,30^. We combined this staining with an anti-mNeon and a nuclear DNA marker, using ultrastructure expansion microscopy (U-ExM) protocol, to achieve higher resolution with greater detail (Fig. 3C). While the exact molecular functions of the *Plasmodium* centrins are somewhat unclear, they are hypothesized to support mitosis by connecting microtubule organization center with the nuclear envelope^30^. It was therefore reassuring to observe that during Branching Phase I and II (ten to four hours before egress) one or two centrin foci decorate each of the dividing nuclei (Fig. 3C). As expected, the fully-branched apicoplast was not stretched yet over nuclei, nor was it found at the vicinity of centrin foci (Fig. 3C). Unfortunately, the fixed nature of this experiment deprived us from the quick Crown-stage parasites. However, subcellular organization of apicoplast-divided parasites (step 4) was highly illuminating (Fig. 3D). In these cells, the already-divided minute apicoplast organelles, in unexpected circular ring forms, were paired with individual nuclei while co-localized with centrin foci (Fig. 3D). Collectively, these results support a role for CPs in apicoplast sporting into daughter cells.

Accurate division and sorting are key to apicoplast inheritance. However, this process is not accomplished solely with physical delivery of the organelles into daughter cells. A central component of apicoplast inheritance is the propagation and delivery of its genetic material to the next generation, namely the organelle’s genome.

### The apicoplast carries a haploid genome which is synchronically replicated with the nuclear DNA

The apicoplast carries its own genome which needs to replicate in order to be inherited into daughter cells. Little is known about cellular or genetic functions of organelle genome, and any potential role in organelle biogenesis is yet to be discovered. Moreover, initial genome copy number and the timing of replication during the cell cycle are unknown. To approach this, we first determined organelle genome copy number (i.e. ploidy). To do that, we optimized a quantitative Real-Time PCR (qRT-PCR) approach performed on DNA (rather than transcripts) of all three genomes; nuclear, apicoplast and mitochondrion. The extremely high AT content, particularly in the apicoplast genome which is close to 90%, poses a challenge for accurate quantification. To overcome this, we tested dozens of primer pairs from the nucleus and the apicoplast, and chose for each genome five segments that could be amplified efficiently and consistently (Table 1). We then chose several of these segments, cloned them into plasmids, and tested amplification at known plasmids concentrations in order to verify accuracy and link Cycle threshold (Ct) values to absolute gene copy number (Fig. 4A). We observed that 10-fold plasmid dilutions manifested in a 3.3 change in Ct value (Fig. 4A, left) and 4-fold gDNA dilution showed a 2-cycle gap in Ct values (Fig. 4A, right), confirming the required 2^ transformation for accurate qRT-PCR measurement. Due to the small size of the mitochondrial genome (6kb), only two primer pairs were chosen, both demonstrating high efficiency (Table 1). To determine organelles’ copy numbers, we synchronized and extracted genomic DNA from early-mid stage parasites, prior to the onset of nuclear replication. Surprisingly, we observed comparable Ct values for the nuclear and apicoplast genomes. Because *P. falciparum* is a haploid parasite, we concluded that the early-stage apicoplast of *P. falciparum* carries a single copy of its circular DNA (Fig. 4B and Fig.S4). This finding was in stark contrast to the mitochondrion genome which appears in significantly higher copy numbers, either representing an earlier replication onset, a total higher copy number or a combination of both (Fig. 4B and Fig.S4).

**Table 1.**
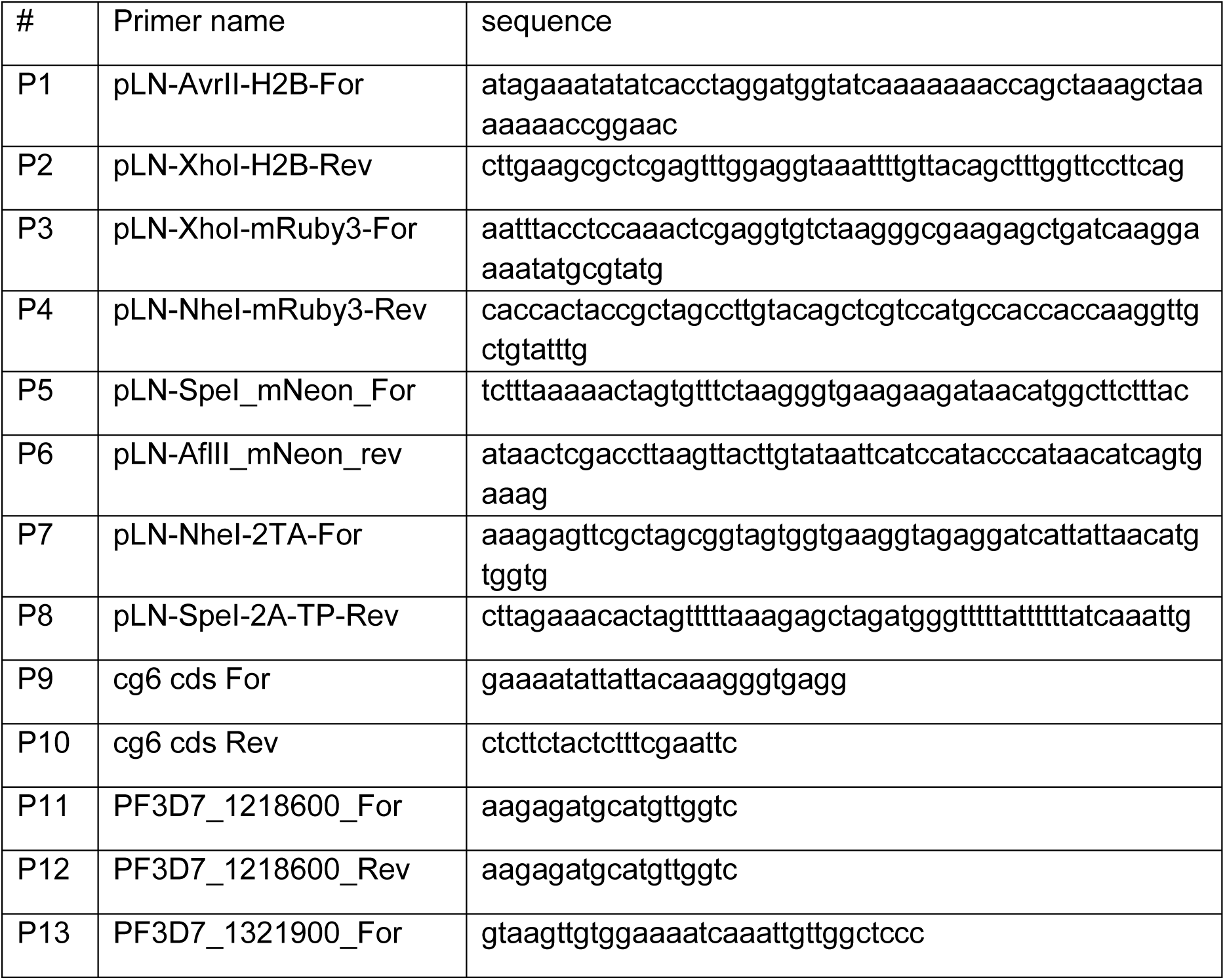
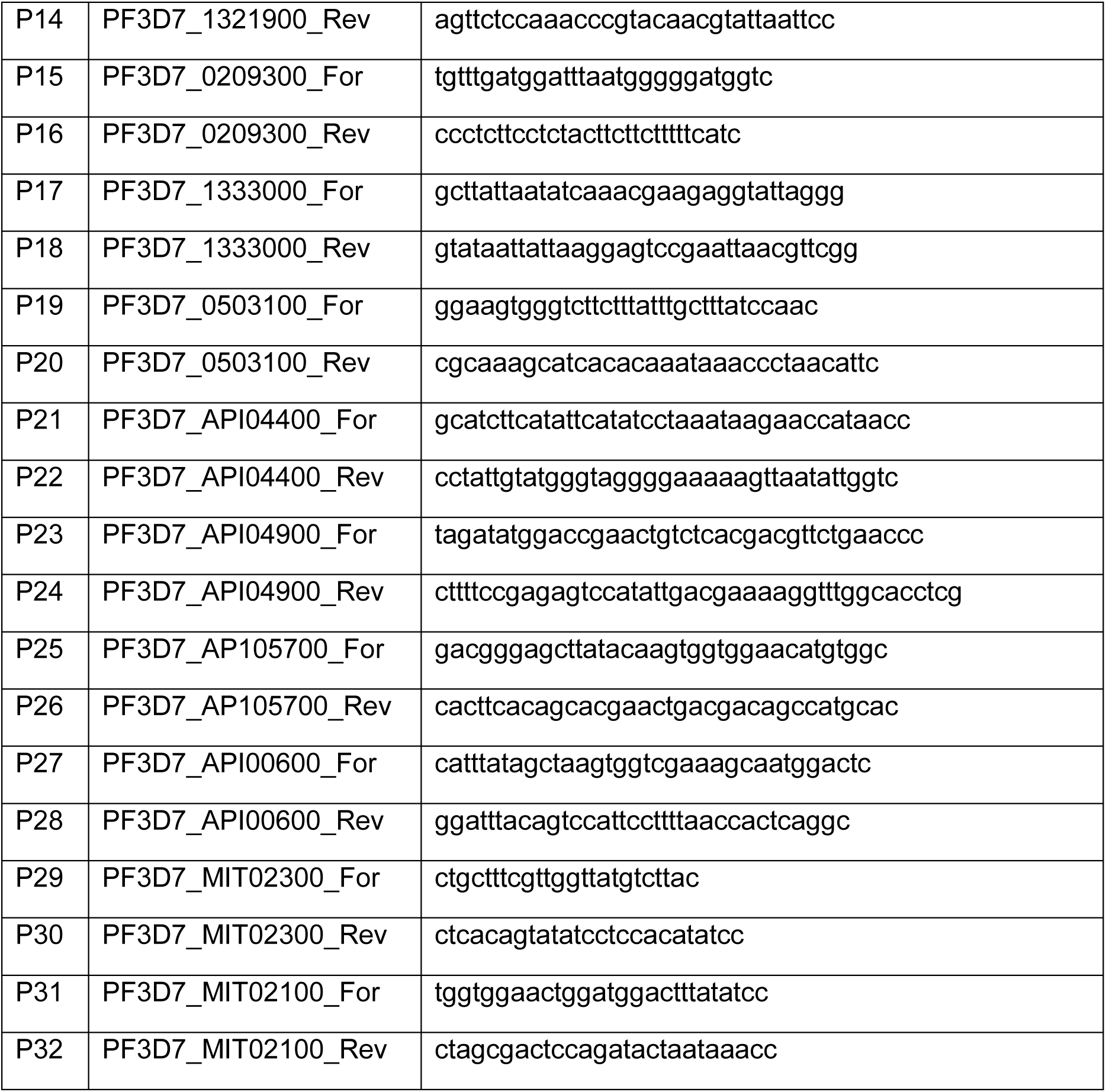

**Figure 4.**
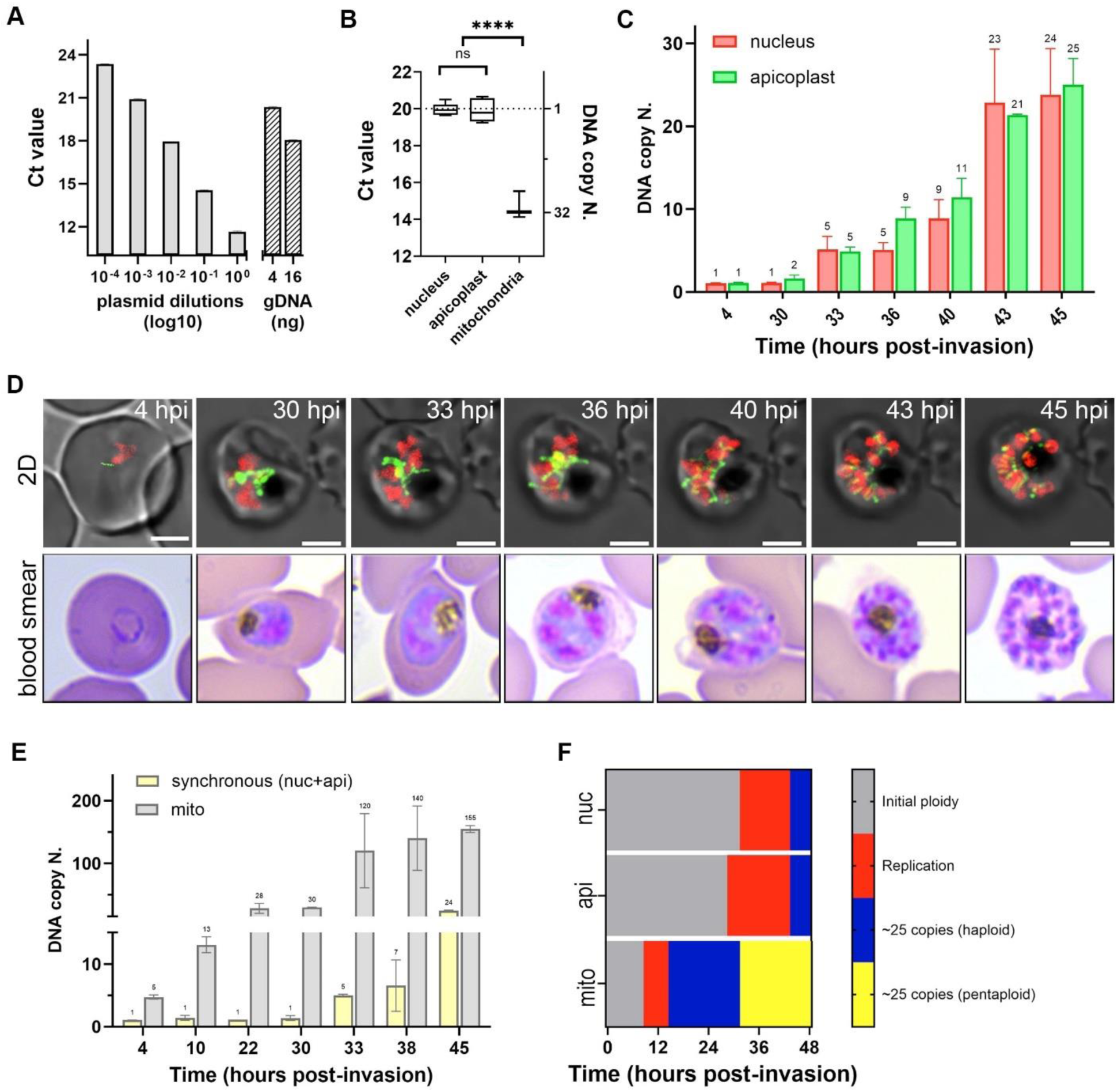
Kinetics and Synchrony of nuclear, apicoplast, and mitochondrial genomes replications throughout the parasites’ cell cycle. **A.** Quantification of gene copy number using plasmid standards and genomic DNA (gDNA) via quantitative real-time PCR (qRT-PCR) performed on DNA. A nuclear gene (PF3D7_1321900) was amplified to assess primer efficiency. Plasmid DNA containing the target sequence was serially diluted (1:10) from 1 ng/µL to 0.1 pg/µL, yielding an expected cycle threshold (Ct) shift of ∼3.3 cycles per dilution. In parallel, gDNA was serially diluted (1:4), demonstrating a Ct shift of ∼2 cycles per dilution. Absolute gene copy numbers in gDNA samples were determined by interpolating Ct values against the plasmid standard curve. The x-axis represents plasmid dilutions in log10 scale and gDNA dilutions in ng, while the y-axis denotes Ct values. **B.** Comparative qRT-PCR analysis of nuclear, apicoplast, and mitochondrial genomes in early ring-stage *P. falciparum* parasites. DNA was extracted, and qRT-PCR was performed using primers targeting representative genes from each genome. For each primer set, 3 technical replicates from 3 independent experiments were used. To determine ploidy for each genome, values from two-five primers sets (i.e. segments per genome) were averaged. Calculated Ct values reveal that nuclear and apicoplast genomes have on copy number (haploid), whereas mitochondrial genes exhibited lower Ct values, indicating a higher initial genome copy number or an earlier replication time. The x-axis represents analyzed genomes, and the y-axis denotes Ct values. **C.** Temporal analysis of nuclear and apicoplast’ genome replication during the intraerythrocytic cycle. qRT-PCR was performed on synchronized parasite cultures sampled at various time points over the 48-hour intraerythrocytic life cycle. Both nuclear and apicoplast’ genomes initiate replication at ∼33 hours post-invasion, with genome copy numbers increasing to an average of ∼25 per schizont, corresponding to daughter cell formation. The x-axis represents different time points in the asexual cycle, while the y-axis denotes genome copy number. **D.** Representative long-term fluorescence live-cell microscopy images of the *P. falciparum* reporter line H2B-mRuby/TP-mNeon (upper panel), complemented with Giemsa-stained blood smears (lower panel) for morphological assessments at key time points. **E.** Mitochondrial genome replication dynamics (grey) compared to nuclear and apicoplast genome synchronous replication (yellow) during the *P. falciparum* intraerythrocytic cycle. qRT-PCR analysis of DNA from synchronized parasites revealed that the mitochondrial genome starts with approximately five copies per parasite. Replication initiates ∼10 hours post-invasion and is completed by 33 hours post-invasion, reaching 100–150 copies per schizont. This corresponds to ∼5 times the haploid genome copy number observed for nuclear and apicoplast genomes (mean of both, yellow). The x-axis represents time points over the 48-hour cycle, while the y-axis denotes mitochondrial genome copy number. **F.** Temporal mapping of nuclear (nuc), apicoplast (api), and mitochondrial (mito) genome replication throughout the intraerythrocytic cycle. The mitochondrial genome (red) undergoes replication first and reaches its final copy number (yellow) before nuclear genome replication begins. In contrast, nuclear and apicoplast genome replication (red) occurs synchronously at a later stage, reaching the final haploid genome copy number ∼3 hours before egress (blue). The x-axis represents hours post-invasion, while the y-axis displays the three organellar genomes analyzed.

Next, we measured DNA amplification timing and rate during the life cycle. Tightly synchronous H2B-mRuby-2TA-TP-mNeon parasites were grown and monitored for the entirety of the 48-hour cell cycle. Every two hours, samples were taken for: (1) genomic DNA extraction to calculate copy number, (2) blood smear to follow overall parasite stage, and (3) fluorescent live imaging to follow nuclear and apicoplast development (Fig. 4C and 4D). We found that replications of both genomes, nuclear and apicoplast, begin simultaneously around 30 hours post invasion (Fig. 4C), at the time point in which we observed the onset of apicoplast’s morphological changes (Fig. 4D). Despite the fact that organelle and nuclear genomes highly differ in size and replicate via two distinct replication machineries, we found a perfect synchrony at their replication rate. In each time point measured, their average copy number was identical (Fig. 4C). Both reached a final number of ∼25 DNA copies (nucleus 24±6; apicoplast 25±3) three hours before egress at the Crown stage, prior to organelle division and cytokinesis. This again was in a striking contrast to mitochondrial genome, which began with an initial number of five copies (hence, a pentaploid) (Fig. 4E). The mitochondrion also amplifies its genome copies by approximately 25-fold. However, this occurs much earlier in the cell cycle, beginning roughly 10 hours post invasion at the early ring stage (Fig. 4E). Consequently, mitochondrial genome replication is fully completed with ∼25 sets of pentaploid genomes by the time nuclear and apicoplast replications begin 33 hours post invasion (Fig. 4F).

### Inhibition of cell cycle S-phase reveals dependency between apicoplast biogenesis and nuclear replication

The spatial and temporal coordination between nuclei divisions and apicoplast morphogenesis suggest a central regulatory mechanism. Lacking known factors driving apicoplast biogenesis throughout the cell cycle, we hypothesized that DNA replication may play a regulatory function. To explore the role of nuclear genome replication, we used the drug aphidicolin, an inhibitor of eukaryotic DNA polymerase Alpha and Delta^31^. Aphidicolin has been shown to reversibly block *Plasmodium* cell cycle at the early S phase by efficiently suppressing nuclear DNA replication^32,33^. To determine the required dose for aphidicolin treatment, we performed a half-maximal effective concentration assay (EC50), and found it to be 0.7 μM (Fig.S5A). We therefore incubated the parasites with 3X EC50 (2 μM) aphidicolin, a concentration significantly lower than previously used^33^, to minimize toxicity. Following parasites development in the presence of aphidicolin, we observed a block in nuclear division, very rarely proceeding beyond two nuclei (Fig. 5A and Sup. Movie 10). This block was also manifested as stagnation in nuclear volume, most notably in the last ten hours before egress (Fig. 5B and Fig.S5B). Importantly, blocking S-phase initiation inhibited apicoplast development in its initial stages (Fig. 5A). The organelle began Elongation (step 1) but stopped at the first phase of Branching (step 2I), and did not further change morphology (Fig. 5A), volume (Fig. 5C and Fig.S5C) or structure as manifested by surface area: volume ratio (Fig. 5D). Consequently, final total organelle numbers upon treatment (apicoplast and nuclei) were extremely low but equal (Fig. 5E). These results demonstrate tight coordination and dependency between S-phase initiation, nuclear DNA replication and apicoplast biogenesis.

**Figure 5.**
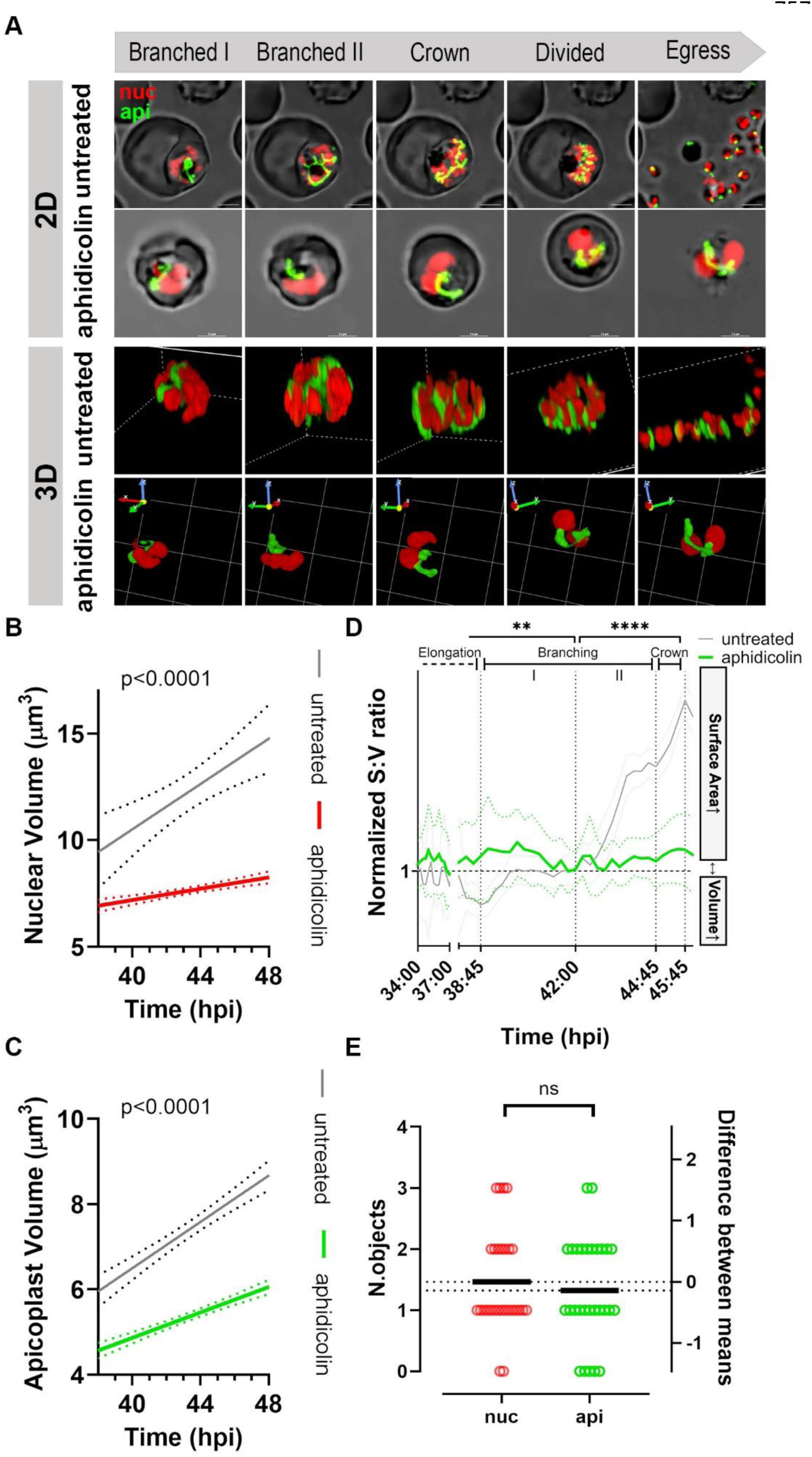
Nuclear DNA replication determines the onset and progression of apicoplast biogenesis. **A.** Representative images from sequential capturing of apicoplast biogenesis and nuclear replication of aphidicolin-treated H2B-mRuby/TP-mNeon *P. falciparum* parasites. Detailed apicoplast (green) and nucleus (red) morphologies and conformations during different stages of parasite’s life cycle in 2D (maximum intensity projection, upper panel) and 3D (volume visualization, lower panel) using NIS image analysis software. In untreated parasites, the nucleus and apicoplast follow divisions and normal biogenesis, whereas in 2μM aphidicolin treated parasites nucleus divisions are stalled and a single undeveloped apicoplast is seen, indicating a link between replication and impairment in organelle biogenesis. 2D Scale bars, 2.5 μm. 3D grid 2×2 μm. **B.** Linear regression analysis shows a significant reduction in nuclear volume at last 10 h of the life cycle with 2μM Aphidicolin treatment, p<0.0001. Aphidicolin-treated red, untreated grey. **C.** Linear regression analysis show significant impairment of apicoplast volume at last 10 h of the life cycle with 2μM aphidicolin treatment, p<0.0001. Aphidicolin-treated green, untreated grey. **D.** Quantification of median apicoplast surface area: volume vs. time was obtained as described in Materials and Methods for n=28 parasites for each time point, from 2 independent experiments. SA: V ratio in aphidicolin-treated parasites (green line) represents mathematically the absence of branching of the organelle at the last 6 h of the cell cycle. Dashed grey line serves as a reference to SA:V ratio in untreated parasites, for which data are derived from Fig. 2F. **E.** Final number of organelles per parasite was calculated as described in Materials and Methods for n=28 parasites from two independent experiments. The difference between nuclei and apicoplast mean numbers (2 (nuc) and 2 (api), (api - nuc) = -0.07143) is not significant, paired t test, P=0.8425.

### Replication of apicoplast genome is required for Crown formation

To investigate the link between replication of apicoplast genome and organelle development, we used ciprofloxacin, an antibiotic that prevents prokaryotic DNA replication through inhibition of DNA gyrase^34^. We determined the half-maximal effective concentration and observed a sharp drop in EC50 value after 96h of drug treatment (Fig.S6A,B,C). This effect is typically observed with apicoplast targeting drugs like doxycycline and chloramphenicol^35^. Like them, ciprofloxacin (CIP) appears to kill parasites on their second replication cycle, in a mode termed delayed cell death. To further verify that CIP activity is specific to the apicoplast, we tested the effect of the drug in the presence of isopentenyl pyrophosphate (IPP), the essential apicoplast-producing metabolite^36^. We observed that CIP+IPP treatment restored parasites growth, verifying apicoplast-specificity (Fig.S6D). Critically, this enables us to uncouple apicoplast genome inhibition from secondary apicoplast-related metabolic defects. Therefore, from this point, these experiments were done under CIP+IPP conditions.

We proceeded to test the effect of CIP treatment during the first replication cycle, before any effect on parasite viability is observed. We incubated parasites with CIP+IPP throughout the entire cycle and monitored nuclear and apicoplast development using dynamic imaging (Fig 6A and Sup. Movie 11). We observed an overall normal cell cycle progression, with small yet significant apicoplast aberrations at the very last stages of the cell cycle (Fig. 6A). As expected, nuclear replication was unaffected, as could be seen in the images (Fig. 6A and Sup. Movie 11) and in nuclear volume quantification (Fig. 6B and Fig.S6E). The effect of CIP+IPP on organellar development, though, was more complex. Parasites did complete the cycle and egressed, however with certain organellar abnormalities (Fig. 6A). Apicoplast volume increased normally until the last five hours before egress and then stopped (Fig. 6C and Fig.S6F). This was mostly apparent in the surface area to volume (SA:V) ratio, where the sharp increase that is expected five hours before egress, disappeared upon CIP+IPP treatment (Fig. 6D). Careful examination revealed that the SA:V ratio dropped only at the last stages because certain morphologies could not be formed (Fig. 6D and 6A). Although the apicoplast elongated (step 1) and branched (step 2) normally during CIP treatment, it did not accomplish the full Crown morphology (step 3). This quick step of stretching a single apicoplast over the final number of nuclei, was the only missing step upon CIP+IPP treatment. The apicoplast then moved to division (step 4) which was completed but with consequences. Under normal conditions the apicoplast forms the Crown, comes into close association with all nuclei via tubulin, divides, and is sorted equally in a 1:1 ratio into daughter cells (Fig.6A, panel 3D, untreated). Upon CIP+IPP treatment, the Crown morphology is missing, but apicoplast biogenesis proceeds by what seems superficially like normal division (Fig.6A, panel 3D, treated). However divided organelles were no longer associated with individual nuclei and deviated from their 1:1 ratio (Fig. 6A and 6E). This indicates that apicoplast genome replication does not control or required for most steps of organelle biogenesis, including Elongation (step 1), Branching (step 2) or Division (step 4). However, it is required for Crown formation (step 3), and its absence leads to failure in apicoplast sorting into daughter cells.

**Figure 6.**
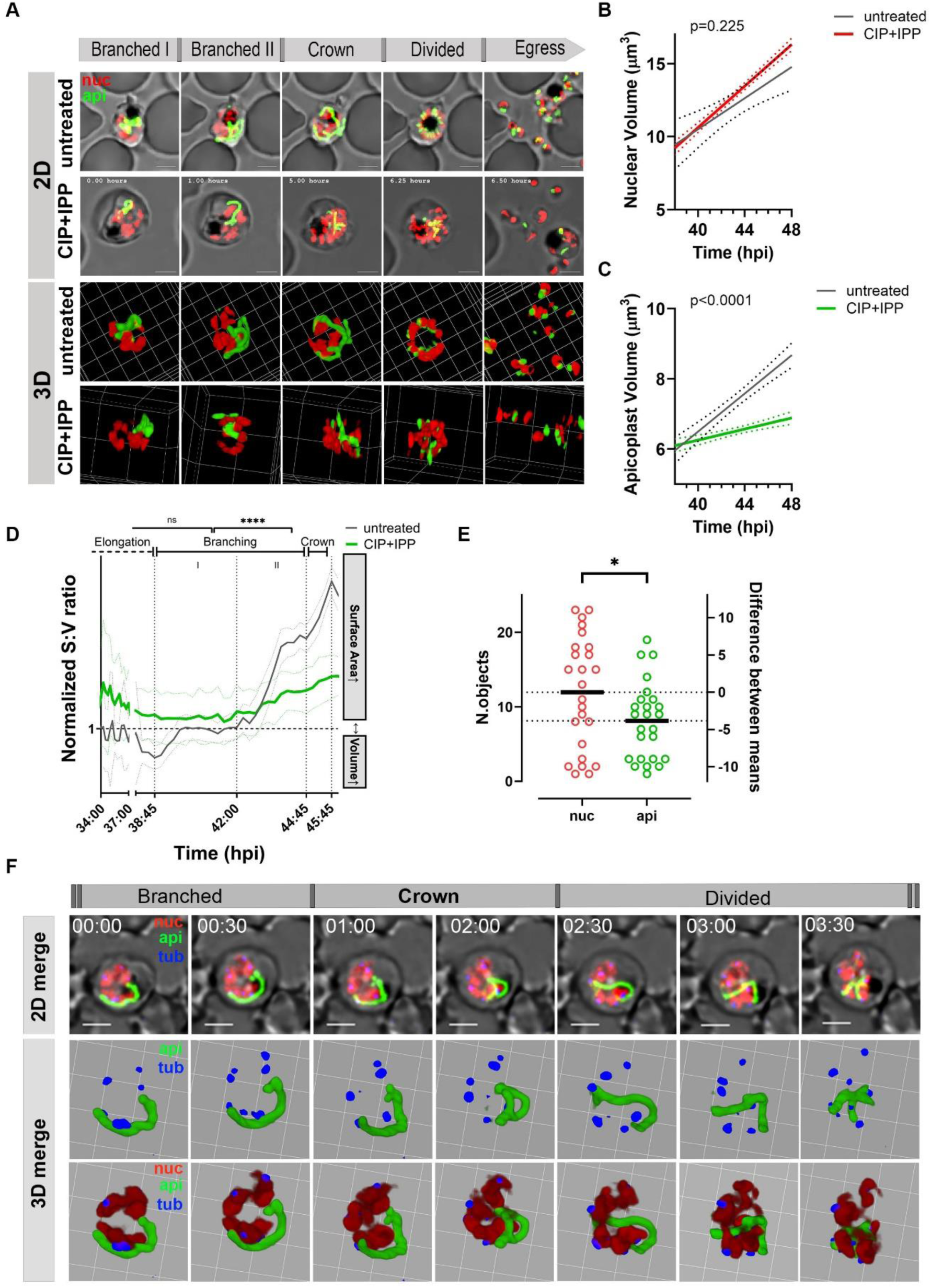
Apicoplast genome replication is requires for Crown morphology formation and accurate sorting into daughter cells. **A.** H2B-mRuby/TP-mNeon *P. falciparum* parasites were treated with 8μM ciprofloxacin (CIP) to inhibit apicoplast DNA gyrase and 200μM isopentenyl pyrophosphate (IPP) to guarantee proper apicoplast metabolism. Representative images from sequential capturing of apicoplast biogenesis and nuclear replication upon CIP+IPP treatment. Detailed apicoplast (green) and nucleus (red) morphologies and conformations during different stages of parasite’s life cycle in 2D (maximum intensity projection, upper panel) and 3D (volume visualization, lower panel) using NIS image analysis software. In untreated parasites, the nucleus and apicoplast follow divisions and normal biogenesis are seen. Whereas in 8μM CIP/200μM IPP treated parasites, the nucleus undergoes multiple divisions, but the apicoplast’s crown structure remains incomplete, and its branches are not observed next to every nucleus in the cell. Scale bars, 2.5 μm. 3D grid in untreated is 2×2 μm and in treated is 5×5 μm. **B.** Linear regression analysis shows no effect on nuclear volume at last 10 h of the life cycle upon CIP+IPP treatment, p=0.225. CIP+IPP-treated graph is red, Untreated is grey. **C.** Linear regression analysis show significant impairment of organellar volume at last 10 h of the life cycle with CIP+IPP treatment, p<0.0001. CIP+IPP-treated graph is green, Untreated is grey. **D.** Quantification of median organelle surface area: volume vs. time. Data were obtained as described in Materials and Methods for n=25 parasites for each time point, from two independent experiments. SA: V ratio in treated parasites (green line) is a mathematical representation of the absence of crown formation at the last 6 h of the cell cycle. grey line serves as a reference to SA:V ratio in untreated parasites, for which data are derived from Fig. 2F. **E.** Final number of organelles per parasite was calculated as described in Materials and Methods for n=25 parasites from two independent experiments. The difference between nuclei and apicoplast mean numbers (12 nuclei and 8.1 apicoplast organelles, (api - nuc = - 3.840), paired t test, P=0.0014) is indicative of daughter cells without an apicoplast. **F.** 8μM CIP + 200μM IPP treatment inhjibits organelle DNA replication and causes impairment of apicoplast-tubulin-nuclei association, leading to nuclei-tubulin structures without adjacent apicoplast, even at late crown/ division stages. 3D grid 2×2 μm.

Lastly, we performed tubulin-tracking live experiments during CIP+IPP treatment. As expected, parasites’ cells developed normally, egressed and reinvaded with the exception of an incomplete Crown step and an aberrant organelle sorting after division (Fig. 6F). Strikingly, tubulin was associated with and orchestrated nuclear divisions, but did not come into close contact with the entire apicoplast as it does in untreated cells (Fig. 6F). Under CIP+IPP treatment, the spatial distribution of nuclei-tubulin in the cell is still predetermined during late Branching (step 2). However, the apicoplast no longer conforms to the nuclei-tubulin-dictating arrangement just before its division. These results suggest that tubulin-apicoplast association during the Crown stage is dependent on organelle genome replication. Furthermore, this association is functionally linked with organelle segregation into daughter cells (see model in Fig. 7).

**Figure 7:**
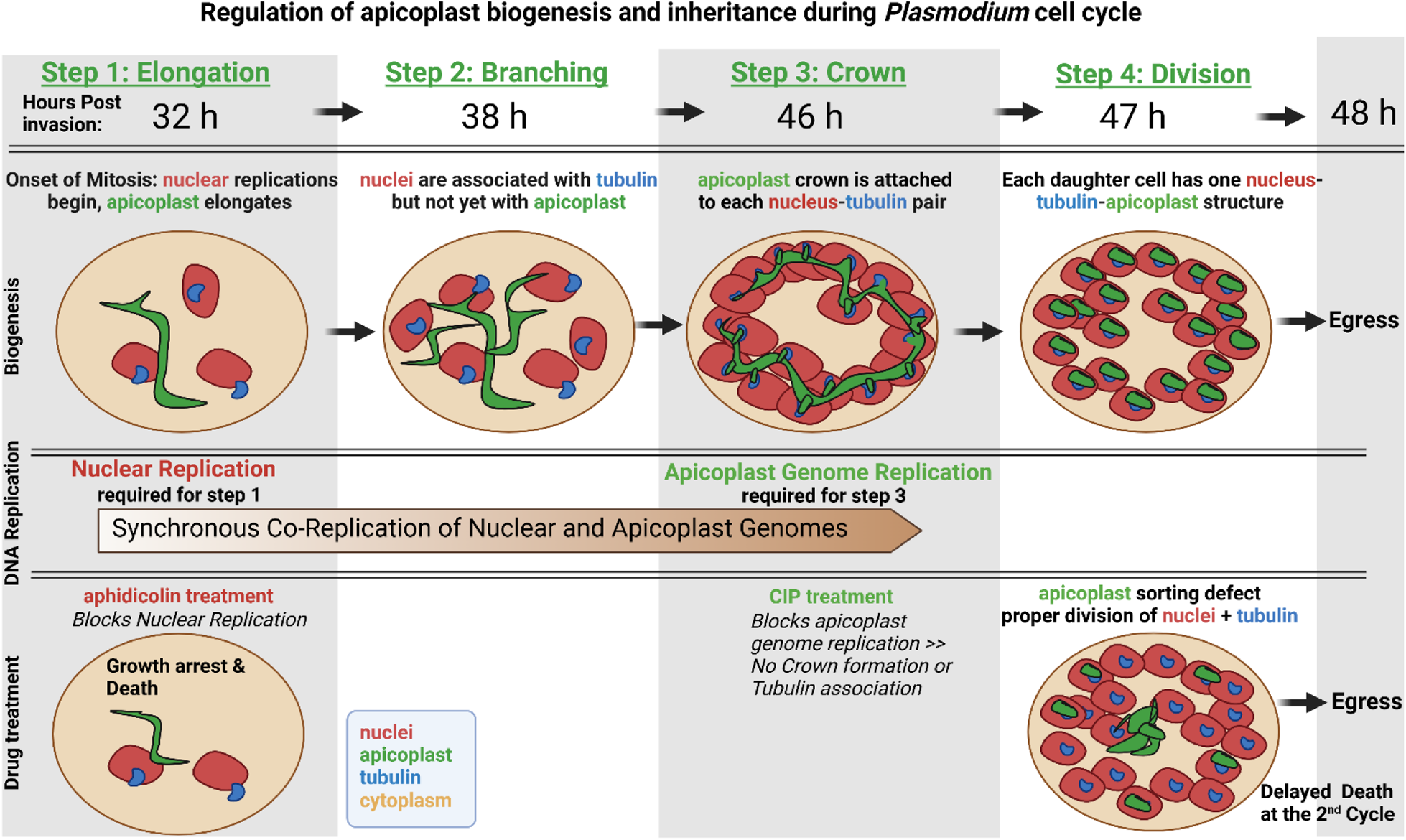
A model for regulation of apicoplast biogenesis and inheritance during *Plasmodium* Intraerythrocytic cell cycle. <u>Step 1 – Elongation:</u> Roughly 32 hours post parasite’s invasion into the host red blood cell, parasite’s mitosis is initiated. When nuclear replication begins (red), the apicoplast (green) begins Elongation, the first step in its biogenesis. These six hours of linear growth coincide with the onset of DNA replications in the nucleus and in the apicoplast. Nuclear and apicoplast DNA replications go hand-by-hand from this point onward, despite being mediated by two distinct replication machineries. Importantly, inhibiting the nuclear DNA polymerase by the drug aphidicolin, also inhibits apicoplast biogenesis, despite not affecting apicoplast genome replication. This suggests a central nuclear regulatory pathway, mediated by a diffusive signal or any other atypical cell cycle checkpoint. <u>Step 2 – Branching:</u> Five to six hours later, the second step in apicoplast biogenesis begins. These six hours of splitting and diverging growing structures are divided into two phases (Branching I and II) based on distinct SA:V ratio signatures. During Elongation (1) and Branching (2), the nuclei and the apicoplast grow at the same pace, almost to the same dimensions, demonstrating tight temporal and spatial synchronizations, despite minimal physical association. <u>Step 3 – Crown:</u> This morphology is characterized by a closed circle with minute budding structures. During the quick one-hour Crown step, the yet undivided apicoplast is stretched over the divided nuclei, adapting to their subcellular distribution. This attachment is mediated via centriolar plaque tubulin and is important for accurate sorting of daughter organelles after division. Critically, when apicoplast genome replication is blocked by the drug ciprofloxacin (CIP), the only step in apicoplast biogenesis being affected is the Crown morphology. This observation suggests that the apicoplast genome plays an essential and unique role in mediating nuclear-tubulin association during the crown stage, and is required for organelle sorting into daughter cells. <u>Step 4 – Division:</u> One hour prior to egress, the apicoplast is divided, and each daughter-organelle is paired with a single nucleus via centriolar plaques. CIP treatment does not directly inhibit this step; however, the absence of the previous Crown formation leads to defective organelle sorting, and to daughter cells lacking an apicoplast.

## Discussion

In this work, we aimed to characterize the details of apicoplast biogenesis during *Plasmodium falciparum* cell cycle. In particular, we were interested in the nature and kinetics of its sequential morphological changes, and in the unknown organizing mechanisms. To do that, we optimized a parasite-adjusted, apicoplast-centered, dynamic imaging platform. Due to the parasite’s minute size and its acute sensitivity to oxidative stress, viable and long-term live-imaging proved challenging to an extreme. Two very recent and related studies overcame these challenges by various methodologies. Klaus *et al* deciphered the asynchronous nature of *Plasmodium* cell cycle by dissecting nuclear replication in high temporal resolution^19^, whereas Morano *et al* revealed a dual role of a dynamin orthologue in organelle fission^18^. Building on and inspired by these works, we added here additional technological advantages that enable us to reconstruct the entire life cycle (48 hours), using high temporal (<15 min) and spatial (<0.5 μm) resolutions. Furthermore, we established here an analytical pipeline that enables us to measure and quantify subcellular dimensions. Using these computational tools, allowed us to test these events in hundreds of cells, extract quantifiable data, and determine their statistical power. Thus, we present here visual and kinetic data of high spatial and temporal resolutions, while being statistically robust.

Unlike a chloroplast, the apicoplast, a product of a secondary endosymbiosis, is ‘twice-removed’ from the cell nucleus. Various previous studies, including our own, pointed to an incomplete communication between the organelle and the nucleus. This is manifested in the ability of the parasite to live well without an apicoplast, as long as essential metabolites are added^36^; the unaltered transcriptomics of nuclear-encoded genes when the apicoplast is removed^23^; and the persistent translation, packaging and trafficking of apicoplast-targeted proteins, even when there is no organelle to dock-in^24,37–39^. This study, however, showed that the apicoplast is by no means an autonomous unit in the cell. Following and measuring both nuclear replication and apicoplast development, we show that the latter is tightly linked and dependent on the cell cycle. The two events were measured by volume, surface area, SA:V ratio and number of objects. Each parameter reveals the same phenomenon, that the nuclei and the apicoplast coordinately act as a single unit during schizogony. This is also interesting because, as Klaus *et al* showed, replication of nuclei is asynchronous and their final number is not predetermined^19^. Most importantly, when we blocked nuclear replication with the drug aphidicolin, it completely blocked apicoplast biogenesis. This is despite the fact that aphidicolin could not directly affect anything in the apicoplast, due to complete absence of any eukaryotic mechanisms in the organelle. Thus, an unknown signal must travel back and forth between the two organelles (and potentially others) to coordinate a highly dynamic event of asynchronous schizogony. In the absence of canonical cell cycle checkpoints in *P. falciparum*^31^, the unknown nature of such signal is intriguing.

Another unexpected observation is that the single apicoplast organelle carries only a single copy of its genome. This organelle haploidy correlates with the haploid nucleus, but deviates from other known endosymbiotic organelles, such as mitochondria and chloroplasts, which carry at least dozens of DNA copies. This is a crucial feature in most organisms, enabling organelle heteroplasmy, the ability to carry genetic polymorphism within a single organelle or a cell. Moreover, organelle genome polyploidy safeguards its genetic material in the case of mutational errors and provides a template to fix them if possible. The apicoplast is thus deprived of all, as well as the opportunity for ‘healthy’ recombination events, that are available to the haploid nucleus during sexual development. Thus, it is very unclear from an evolutionary point of view, how the apicoplast maintains a stable genome; How does the apicoplast avoid accumulating harmful mutations without a second template or recombination, while relying solely on ‘maternal’ inheritance. It will be very interesting to discover the properties of its DNA repair machinery, if one exists. In a strike contrast, the *Plasmodium* mitochondrion carries a pentaploid (five-copies) genome, and exhibits distinct replication kinetics. As we show here, the mitochondrial pentaploid genome is amplified by approximately 25 folds (∼125 copies) at the time that the other genomes only begin replication. The relationship between the number of mitochondrial genomes at the onset of nuclear and apicoplast genome replication is intriguing. The fact that mitochondrial genome has reached at this point to its final replication factor, suggests that the final number of daughter cells has been determined. Future studies are needed to shed light on mitochondrial genome replication rate and a putative role in predetermining the final number of daughter nuclei.

The dynamic and quantitative nature of our studies, enabled us to describe a detailed landscape of apicoplast and nuclear behavior, organelle genomes replications and the coordinated harmony in which all of these progress during the cell cycle. For example, we can now conclude that the onset of schizogony, defined as the first nuclear replication, occurs at 16 hours pre-egress, and dictates the onset of two related, yet independent processes; apicoplast biogenesis and apicoplast genome replication. We also show using various measurements, that the final number of daughter cells, at least for this parasite strain, is between 20-25, significantly lower than the text-book figure of 32 nascent merozoites. Importantly, we defined here four discrete morphological steps in apicoplast development, with one of them (Crown, step 3) described here for the first time. The Crown morphology is a quick, one-hour-long step, in which the yet-undivided organelle becomes associated with each of the newly-divided nuclei. We further showed that this association is mediated via centriolar tubulin that are attached to each nucleus in the multinucleate schizont.

Most illuminating is the role of the apicoplast genome in organelle biogenesis and sorting. Quite surprisingly, ciprofloxacin, which blocks the apicoplast-targeted DNA gyrase, did not affect organelle biogenesis despite blocking organelle genome replication. It did, however, affect one specific step in organelle development; the crown morphology. While CIP+IPP-treated parasites showed all other apicoplast morphologies, a flaw in one quick step led to dire consequences for apicoplast sorting to daughter cells. It is a known phenomenon that apicoplast-targeted antibiotics kill parasites within their second replication cycle. As our data show, CIP falls into this category, and thus can shed light on the reason for the delayed-death effect. Because apicoplast genome replication is not required for most steps of organelle development, it does not affect the first generation of treated parasites. However, it affects inheritance of organelles to daughter cells, and thus, it is the second generation of treated parasites that die. It is interesting that apicoplast sorting depends both on organelle genome and on association with centriolar microtubules. A direct visualization of apicoplast DNA and its localization during the Crown stage will be fundamental for future studies. Another intriguing hypothesis is that the apicoplast genome carries a gene that encodes a Tubulin-associated protein that guarantees its inheritance across generations. The identity and function of such an apicoplast-encoded gene, if exists, remains to be investigated.

To summarize, our study provides new insights into the intricacies of *Plasmodium falciparum* asynchronous cell cycle, which require even replication of its three genomes and coordinated biogenesis of all subcellular organelles. We show that apicoplast development is tightly coupled with nuclear replication, challenging the idea of apicoplast autonomy. The discovery that the apicoplast relies on nuclear replication for its biogenesis opens the door to new potential drug targets, as targeting the signaling mechanisms between the nucleus and apicoplast will disrupt parasite growth. Its unique features of genome replication and haploidy discriminate the apicoplast from other endosymbionts. Our findings give rise to many new scientific questions, among which the preservation of an intact genome without a second genetic template or sexual recombination. Furthermore, our discovery of specific morphological steps and the role of the apicoplast genome in sorting and inheritance presents new avenues for therapeutic exploration. Given the critical role of bacteria-derived organelles in participating and regulating the cell cycle of malaria parasites, further studies may reveal promising and safe drug interventions.

## Materials and methods

### Plasmids construction and transfections

Genomic DNA was isolated from *P. falciparum* using NucleoSpin Blood kit (Machery-Nagel). All constructs utilized in this study were confirmed by Sanger sequencing (Hylabs). PCR products were inserted into the respective plasmids using In-Fusion Snap assembly Master mix (Takara). All restriction enzymes used in this study were purchased from New England Biolabs. Plasmids were amplified by transformation into 50 μl of competent Stellar *E. coli* cells (New England Biolabs GmbH), overnight culture at 37°C, and extraction via the NucleoBond Xtra Midi Kit (Macherey-Nagel).

Oligonucleotides used in this study are summarized in Table 1. For the generation of H2B-mRuby-2TA-TP-mNeon construct, H2B orthologue (PF3D7_1105100) was amplified from genomic DNA using primers P1+P2. P3+P4 were used to amplify mRuby, P5+P6 were used to amplify mNeon, and P7+P8 were used to amplify T2A (skip peptide). All 4 PCR products were then conjugated together using overlap extension PCR and inserted as one piece between AvrII and AflII into pLN plasmid. The cassette was placed under the control of the cam promoter and the plasmid was co-transfected with a pINT plasmid expressing Bxb1 transposase^25^ for stable insertion into the attB site on chromosome 6 in NF54^attB^ parasites.

Parasites’ transformation was carried out as described before^40^. Briefly, 400 μl packed erythrocytes were electroporated (0.32 kV, 925 μF; Gene Pulser II, Bio-Rad) together with a mixture of 15/30 μg of purified plasmid DNA resuspended in 380 μl of CytoMix solution [120 mM KCl, 0.15 M CaCl2, 2 mM EGTA, 5 mM MgCl2, 10 mM K2HPO4/KH2PO4, and 25 mM Hepes (pH 7.6)]. Schizont-stage parasites were added, and selection was applied with 2.5 µg/mL blasticidin-S (Fermentek) 48-h post-transfection. Primers P9+P10 were used to test for accurate integration in the parasite following transfection. For each transfection, at least 2 clones were isolated via limiting dilutions and used in all subsequent experiments. *Plasmodium falciparum* parasites were maintained in RPMI medium supplemented with Albumax I (Gibco) under standard culturing conditions.

### Parasite Culturing and Synchronization

Parasites were cultured in RPMI medium supplemented with Albumax I (Gibco) and transfected as described earlier^41–43^. Synchronization of cultures were achieved by intermittent sorbitol and Percoll treatments.

### Half-maximal effective concentration (EC50) and Growth assays

EC50 values for aphidicolin and ciprofloxacin were determined by incubating synchronous parasite cultures with serial dilutions (50-0.5μM) of the drugs in 96-well plate. For growth experiments, parasites were sub-cultured to maintain parasitemia between 1 to 4%, while parasitemia was monitored every 24 hours by flow cytometry (Cytoflex) following staining with Hoechst 33342 (Thermo Fisher Scientific). Cumulative parasitemia at each time point was back calculated based on actual parasitemia multiplied by the relevant dilution factors. Data from technical replicates were analyzed and fit to a dose-response equation using Prism (GraphPad Software, Inc.). All experiments shown are representative of at least three biological replicates. For IPP rescue, the parasites were incubated with 3xEC50 CIP with or without 200 μM IPP (Isoprenoids LC).

### Dynamic Imaging: Sample preparation

For live-cell imaging, *P. falciparum* parasites were seeded onto sterile 35-mm glass-bottom high dishes (ibidi) coated with Concanavalin A (5 mg/ml; Invitrogen) and rinsed with PBS^44^. Parasite cultures at 1–2% parasitemia were washed twice in incomplete RPMI by centrifugation (1000×g, 30–60 s), then allowed to settle for 10 min at 37°C. Unbound RBCs were removed by washing with incomplete RPMI until a single-cell layer remained. The medium was then replaced with phenol red–free complete RPMI imaging medium)RPMI 1640 with l-glutamine (Sigma), 0.5% AlbuMAX, 0.2 mM hypoxanthine, 25 mM HEPES (pH 7.3), 0.5 mM Trolox, and 12.5 μg/ml gentamicin(, pre-equilibrated to incubator gas conditions for at least 1 hour.

### Live long-term Microscopy

High-resolution images were acquired using a Nikon ECLIPSE Ti2 confocal fluorescence microscope equipped with an Orca FusionBT monochrome camera (Hamamatsu) and a CSU-W1 Spinning Disk unit with a 50 µm pinhole (Yokogawa). The system was integrated with 405/488/561/638 nm LH+ lasers and a CFI Plan Apochromat Lambda D 100× Oil N.A. 1.45, W.D. 0.13 mm objective. Sample temperature and gas conditions were maintained using a Bold Line Gas and Temperature Controller (OKOlab) at 36°C in a humidified chamber with 5% CO₂ and 5% O₂.

For 3D imaging, multichannel images were acquired sequentially using excitation at the appropriate wavelengths with matching emission filters, alongside differential interference contrast (DIC) imaging. Z-stack images (8–10 µm range) were collected at 0.5 µm intervals relative to the home focus plane, across multiple positions using an automated stage and the Perfect Focus System (PFS) for focus stabilization. The time resolution per stack ranged from 15 to 30 minutes.

Images were processed using the Maximum intensity projection or Volume View Extension module of the NIS-Elements AR software package (Nikon). Laser power and exposure time were kept consistent across all samples and experimental replicates. Image processing, analysis, and visualization were performed using NIS-Elements AR, Imaris (Oxford), and Adobe Photoshop. Equivalent brightness and contrast adjustments were applied within each sample group for display purposes.

### Nuclear and Tubulin staining

For nuclear staining, parasites were incubated with SiR-DNA (3 μM; Spirochrome) in phenol red–free complete RPMI imaging medium for 2–5 min before visualization.

Tubulin TrackerTM Deep Red (Invitrogen) at 100nM (without probenecid) was added to Concanavalin A (Invitrogen) immobilized parasites for 1 h incubation in imaging medium. The stained parasites were washed and imaged as described above.

### Image Analysis

Nucleus and the apicoplast parameters in individual parasites were measured by live-cell fluorescence microscopy using mRuby/mNeon signal intensities. The time of egress was manually detected for each cell and served to align the measurements on time-scale of biogenesis. Volume, surface area and number of objects were quantified from the z-stack time-lapse images with two fluorescence channels. Infected RBCs were detected using Fiji and Bio-Formats^45,46^ with maximum intensity projection of the fluorescent channels over the Z- and time-dimensions followed by thresholding. Raw images were cropped around each single infected RBC. Measurements were taken in each cropped image of a single RBC using Fiji^45^ and the ImageJ 3D Suite V4.1.7 plugin^27^. Apicoplast and nucleus objects were segmented using a thresholding. For each infected RBC the following measurements were taken per channel (apicoplast/nucleus) and per time-point: number of 3D objects (connected components), total volume, and surface area. A Google Colab Jupyter Notebook^47^ was used to merge the measurements files, and further analysis was performed using Microsoft Excel and GraphPad Prism version 10.4.1 for Windows.

### Measuring copy number and replication rate of organelle genomes

#### Primer Optimization

Optimized primer sets for improved efficiency were selected, based on GC content higher than 35%. Primer sets with GC content below 30% failed to yield reliable amplification. Primers were designed with lengths ranging from 18 to 42 base pairs, generating qPCR products between 116 and 160 base pairs. Multiple primer pairs were tested across nuclear, apicoplast, and mitochondrial genomes. Five efficient primer pairs were selected for nuclear genome (P11-20), four for apicoplast genome (P21-28), and two primer pairs were chosen for the mitochondrial genome due to its compact size (P29-32). To establish accurate quantification and correlate cycle threshold (Ct) values with absolute gene copy numbers, selected amplicons were cloned into plasmids and used as quantitative standards across a range of known concentrations.

### Genome replication rate: Sample Collection

Synchronized *P. falciparum* cultures were maintained in 150 mL flasks, and experiments were initiated at the early ring stage 4-h post invasion. Parasites were monitored throughout the entire 48-hour asexual blood-stage cycle. 2 mL culture samples for gDNA extraction were collected at regular intervals. gDNA was extracted from collected samples using the Macherey-Nagel DNA extraction kit, and DNA concentration and purity were assessed using a spectrophotometer. Additional 50 μl were taken for thin blood smears and Giemsa stained, and live imaging was performed using spinning disc confocal microscopy as described above.

### Quantitative Real-Time PCR (qRT-PCR)

qRT-PCR was performed on extracted DNA (NucleoSpin Blood kit,Machery-Nagel) using primers targeting nuclear, apicoplast, and mitochondrial genes. Reactions were prepared using the Myra automated liquid handling system in a final reaction volume of 20 µL, containing 10 µL of Luna Universal qPCR Master Mix, 1 µL of DNA sample, 0.5 µL each of forward and reverse primers (10 µM), and 8 µL of ultrapure water. Reactions were conducted using a Quant Studio real-time PCR system (Thermo-Fischer). qPCR data were analyzed using the ΔΔCt method to compare the relative abundance of apicoplast, mitochondrial, and nuclear DNA throughout the parasite’s asexual blood-stage cycle.

### Ultrastructure Expansion Microscopy

Ultrastructure expansion microscopy (UExM) was performed as previously described^14^ with minor modifications. Briefly, 1ml of parasite culture was adhered to poly-d-lysine coated 12- mm coverslips, washed and fixated in 4% PFA for 20 min at 37°C. Following fixation, coverslips were placed cell-side down on gels, incubated at 37°C for 30 min, and transferred to denaturation buffer. Gels were incubated at 95°C for 90 min, then expanded by washing in doubled-distilled water (ddw, MilliQ). The unstained gels were frozen in propyl gallate solution, followed by three 20 min washes in ddw. The thawed gels were then shrunk with five 20 min PBS washes and subsequently blocked (3% BSA) and incubated with primary antibodies overnight. The following day, gels were washed and incubated with secondary antibodies for 2.5 hr. Gels were again washed before re-expansion with three 30 min ddw washes. After blocking, all steps were carried out protecting the sample from light. Stains and antibodies used in this study: mNeonGreen Tag Antibody (1:100, Cell Signaling Technology), anti-centrin, monoclonal 20H5 (1:200, Sigma-Aldrich), Alexa Fluor® 488 goat anti-rabbit (1:250, Cell Signaling Technology), Alexa Fluor® 594 goat anti-mouse (1:100, Cell Signaling Technology), NHS ester Alexa Fluor 405 (1:250, Thermo Fisher Scientific) and SYTOX Deep Red (1:1000, Thermo Fisher Scientific).

### Airyscan imaging

High-resolution images were acquired using the Airyscan detector of a Zeiss LSM880 confocal microscope using the super-resolution mode (SR) and default image processing parameters. The 405, 488, 561 and 640 laser lines were used, and the emitted light was recorded using following filter protocols (BP495-550/LP570 and BP420-480+495-550) and alternating Z-stacks between wavelengths. Z-stacks were recorded at the 0.13 µm intervals.

### Data Availability

All data are included in the manuscript, figures, supplemental figures and movies

## Supporting information

Sup Movies links

## Acknowledgements

We thank Josh Beck for pLN and pINT plasmids; Sabrina Absalon for Expansion microscopy protocol, discussion and advice; Markus Ganter for advice on live imaging; Sigal Ben-Yehuda and William Sullivan for critical reading and comments on the manuscript; Yael Feinstein-Rotkopf at the Imaging Core Facility at the Faculty of Medicine, En Kerem; Ran Ezer at Agenetek for spinning-disk microscopy technical assistance. VEuPathDB and PlasmoDB for scientific, academic and community service. This work was supported by grants from the Israel Science Foundation [400/22 (personal grant) and 2786/22 (equipment grant)] and HUJI start-up funds to AF. AQ is supported by the Carole and Andrew Harper Diversity PhD program, and AT is supported by a PhD scholarship from Science Training Encouraging Peace (STEP) and the Isadore Sharp Einstein Scholarship. AF is supported by the Abisch-Frenkel Faculty Development Lectureship. AF and MS acknowledge support by The Kuvin Center for the Study of Infectious and Tropical Diseases.

## Author Contributions

Conceived and designed the experiments: AF, MS, AQ

Performed the experiments: MS, AQ, EF, AT

Analyzed the data: AF, MS, AQ

Code Writing: MS, YM

Wrote the paper: AF, MS, AQ

### Supplemental information

Sup Figures: Figures S1–S5, Sup Movies1-11.

### Preprint Server

BioRxiv

---

**Figure S1:**
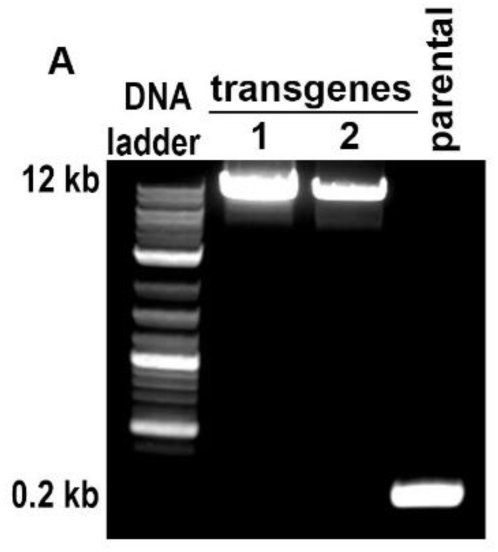
Genotyping of H2B-mRuby/TP-mNeon transgenic parasite line. PCR amplification of genomic DNA isolated from H2B-mRuby-2TA-TP-mNeon transgenic parasite clones. Primers used are (P9+P10) depicted in Table 1. Band size shift from 0.2 to 12 KB appears when integration occurs at the desired attB genomic locus.

**Fig. S2.**
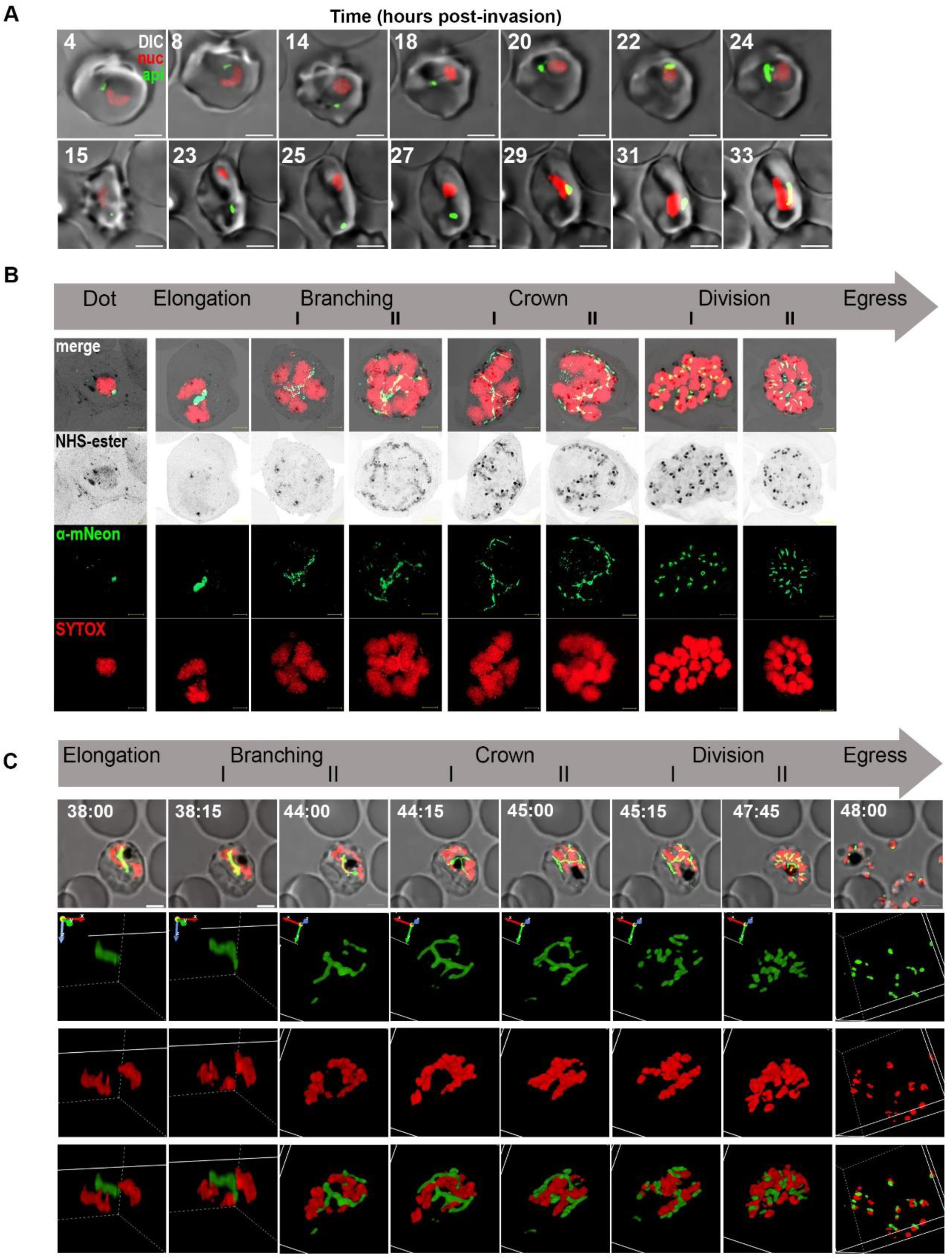
Dynamic and Fixed imaging reveal sequential co-development of nuclei and apicoplast. **A.** Representative images from sequential capturing of apicoplast biogenesis during the first half of the cycle (upper cell) and from late rings stage until late trophozoite stage (15-33 hpi, lower cell). During this time and until schizogony begins roughly 33 hpi, there are no detectable morphological changes to the apicoplast. Images were captured as Z-stacks (total 21 slices of 0.5 μm each) at 100x using spinning disc confocal microscope. The images are displayed as maximum intensity projections (2D) or volume reconstructed for 3D visualization using NIS image analysis software. Scale bar is 2.5 μm. **B.** Representative images of expanded parasites from different cell-cycle stages confirm apicoplast and nuclear morphologies shown by long-term fluorescence live-cell imaging using spinning disc confocal microscope. Apicoplast was stained using anti-mNeon antibody (green), the nucleus was stained by SYTOX Deep Red (red) and 405-conjugated NHS ester stain used for cellular context visualization by labeling protein densities (grey). Images were captured as Z-stacks (10-21 slices of 0.13 μm each) at 63x using Airyscan microscopy. The images are displayed as maximum intensity projections. Scale bar is 5 μm. **C.** Representative long-term fluorescence live-cell microscopy images showing detailed apicoplast (green) and nucleus (red) morphologies and conformations during different stages of parasite’s life cycle in 2D (as maximum intensity projections of Z-stack images) and 3D (volume visualizations). Microscopy Technique: Spinning disk confocal microscopy. Images were captured at 100x as Z-stacks (total 21 slices of 0.5 μm each) with 15 min time intervals for 20 h. Scale bars, 2.5 μm.

**Fig.S3.**
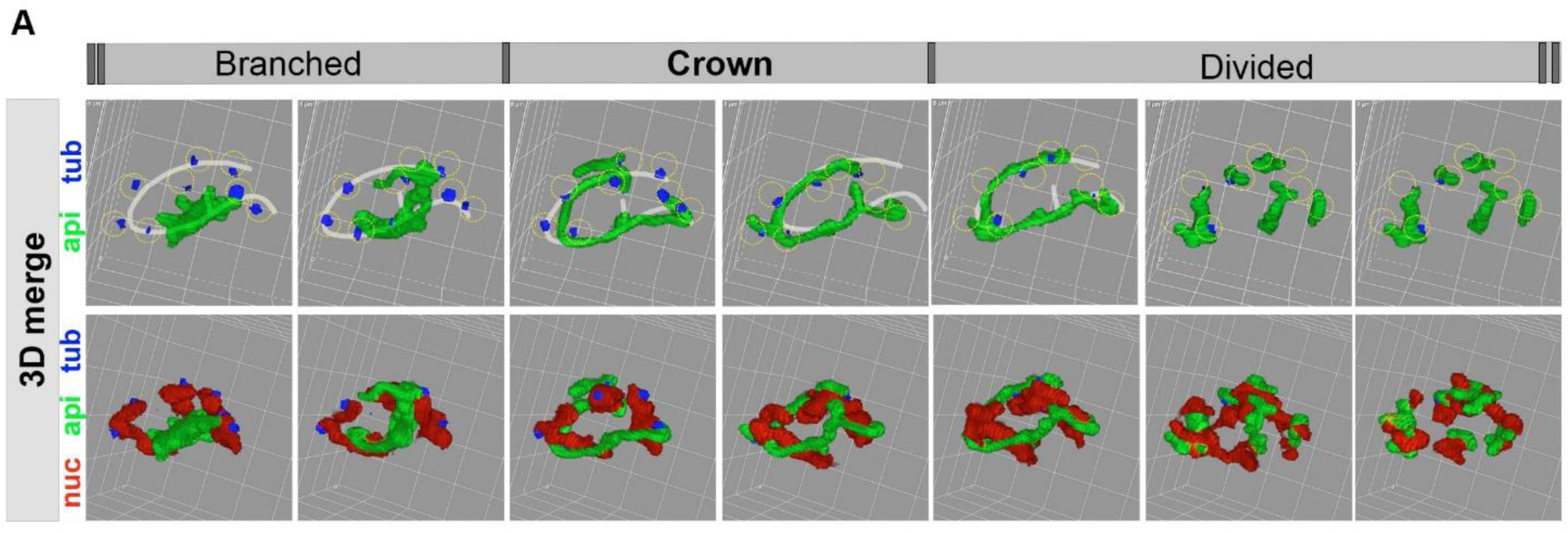
Apicoplast Crown morphology is predetermined by the prior subcellular arrangement and numbers of nuclei-tubulin pairs. Image from Fig. 3A (volume visualization with blue Tubulin Tracker, green apicoplast and red nucleus) marked with final Crown subcellular localization as a white circular line. The white line completely aligns with tubulin-nuclei structures hours before the apicoplast gains its final Crown morphology. This demonstrates that apicoplast Crown morphology is determined by the prior arrangement and numbers of nuclei-tubulin pairs. 3D grid 2×2 μm.

**Fig. S4.**
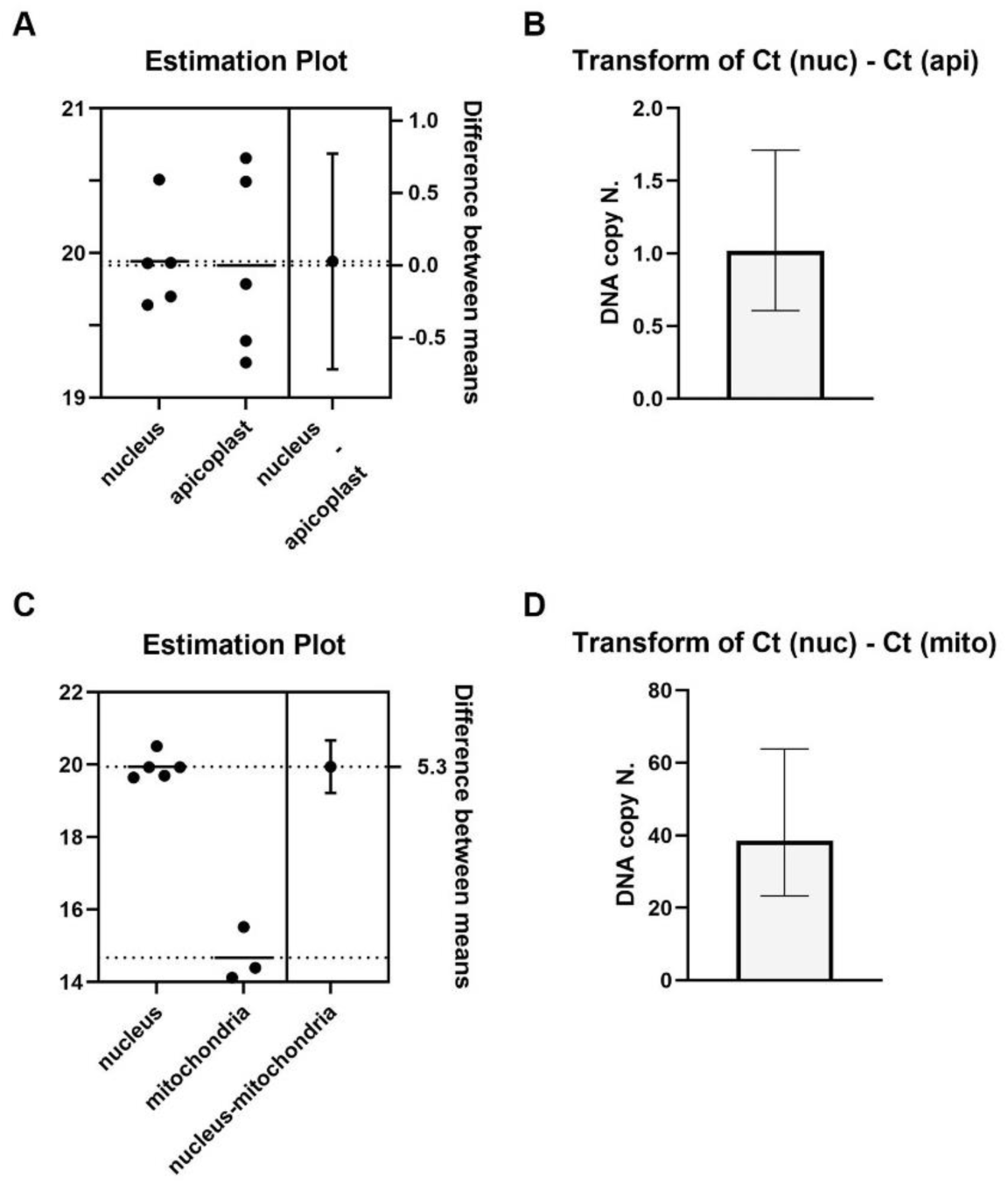
Copy numbers of apicoplast and mitochondrial genomes. **A-B**. Comparative qRT-PCR analysis of nuclear vs. apicoplast (**A**) and mitochondrial (**B**) genomes in early ring-stage P. falciparum parasites. DNA was extracted, and qRT-PCR was performed using primers targeting representative genes from each genome. Five primer sets were used for nuclear and apicoplast genomes, while two sets were used for the mitochondrial genome. Cycle threshold (Ct) values for each genome were plotted as estimation plots, showing Ct values per primer set (mean of triplicates) along with the overall mean Ct for each genome (left Y-axis): nucleus, 19.9 ± 0.3 (**A, B**); apicoplast, 19.9 ± 0.6 (**A**); mitochondrion, 14.7 ± 0.7 (**B**). Differences between means (ΔCt) were analyzed using an unpaired t-test (right Y-axis): Apicoplast vs. nucleus (**A**): difference = 0.028 ± 0.32, P = 0.93 (not significant). Mitochondrion vs. nucleus (**B**): difference = 5.3 ± 0.37, P < 0.0001. **C-D**. To determine ploidy for each genome, the fold difference in genome copy number was calculated as 2^(ΔCt), where ΔCt represents the Ct difference between genomes: 1.019 (- 0.4/+0.7) copies for the apicoplast (C) and 38.56 (-15.3/+25.3) copies for the mitochondria. The calculated genome copy numbers reveal that the nuclear and apicoplast genomes are haploid, while the mitochondrial genome exhibits a higher initial copy number or undergoes replication earlier in the cell cycle.

**Fig. S5.**
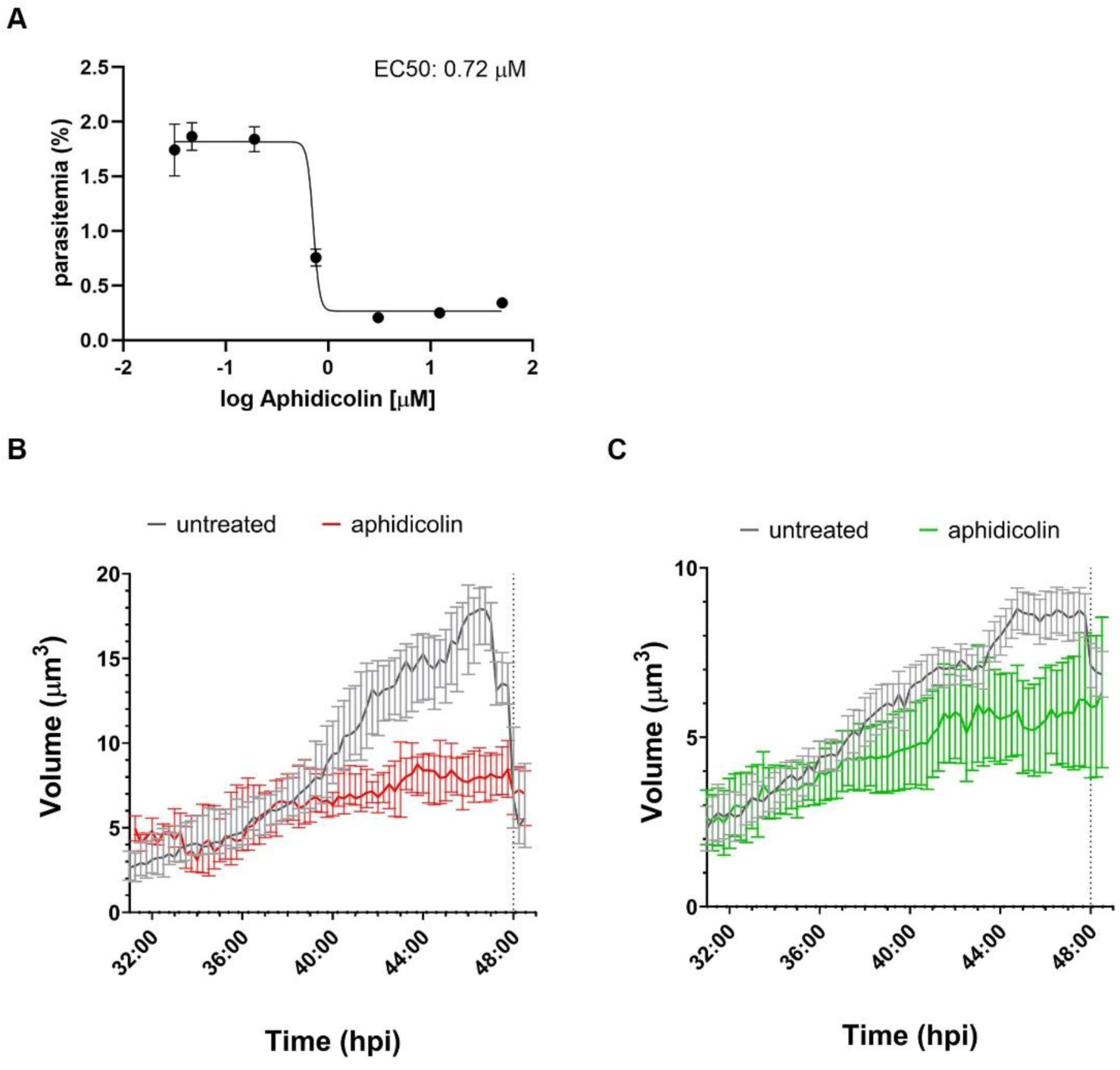
Aphidicolin effect on biogenesis. **A.** To calculate EC50 of aphidicolin, parasites were incubated with serial dilutions of the drug (50-0.05 μm) in a 96-well plate. Parasitemia was measured after 48 h using flow cytometry, showing an EC50 of 0.7 μM. Data are fit to a dose-response equation and are represented as mean ± SEM. One representative experiment out of two is shown. **B.** Quantification of median nuclear volume over time was obtained as described in Materials and Methods for 28 parasites for each time point, from 2 independent experiments. Error bars are 95% CI. **C.** Quantification of median organellar volume over time was obtained as described in Materials and Methods for 28 parasites for each time point, from 2 independent experiments. Error bars are 95% CI.

**Fig. S6.**
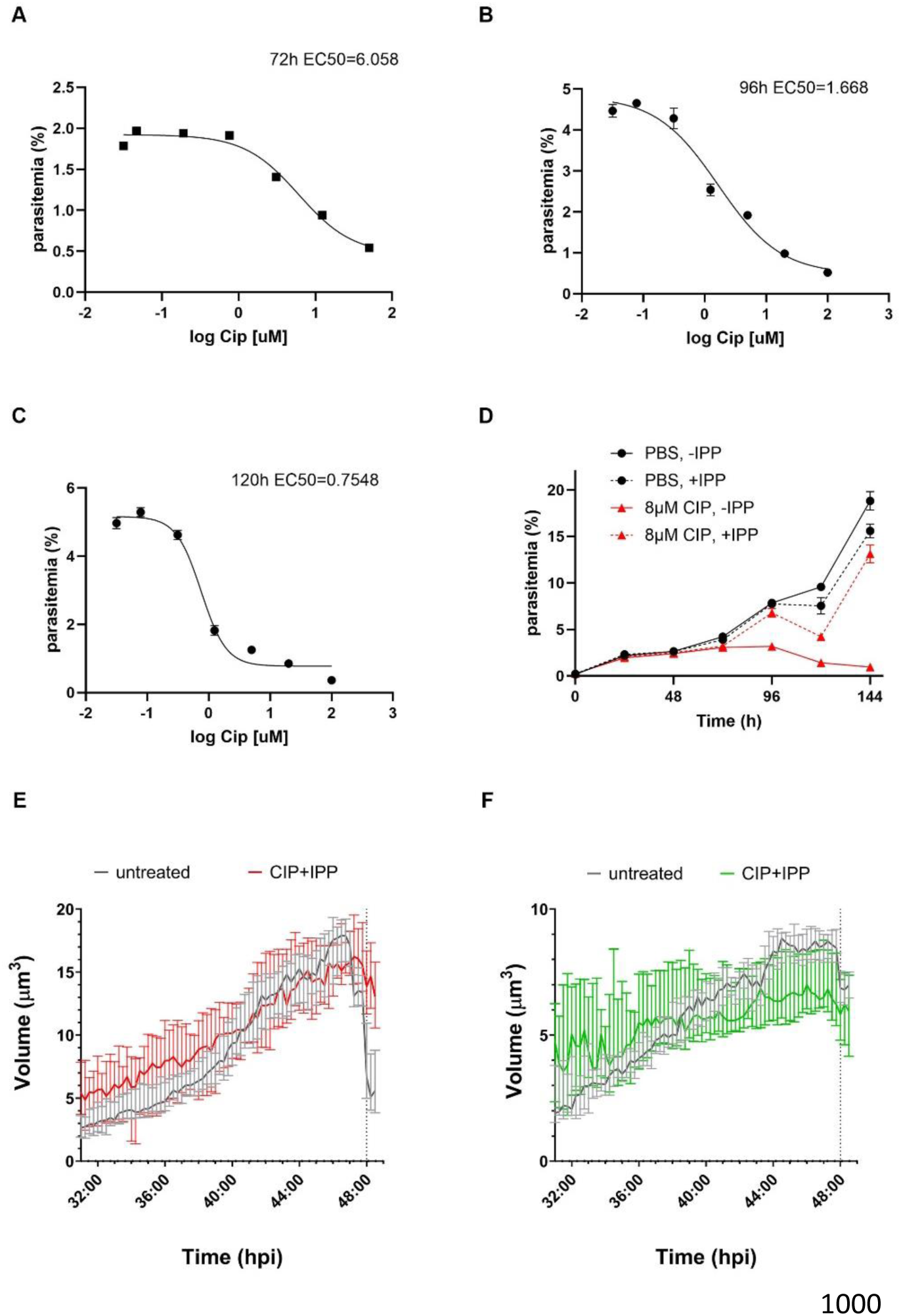
Ciprofloxacin effect on biogenesis. **A-C.** To calculate EC50 of CIP, parasites were incubated with serial dilutions of the drug (50-0.05 μm) in a 96-well plate. Parasitemia was measured at 72 h (**A**), 96 h (**B**) and 72 h (**C**) using flow cytometry showing a gradual decrease in EC50 from 6.058μM to 0.7548 μM. Data are fit to a dose-response equation and are represented as mean of triplicates ± SEM. One representative experiment out of three is shown. **D.** Apicoplast-specific effect of 8 μM CIP treatment was estimated by incubating the parasites with or without 200μM IPP. Parasitemia levels were measured daily by flow cytometry for six days. Data shown as mean of triplicates ± SEM. One representative experiment out of three is shown. **E.** Quantification of median nuclear volume over time was obtained as described in Materials and Methods for 25 parasites for each time point, from 2 independent experiments. Error bars are 95% CI. **F.** Quantification of median organellar volume over time was obtained as described in Materials and Methods for 25 parasites for each time point, from 2 independent experiments. Error bars are 95% CI.

**Sup. Movie S1 ([link](https://drive.google.com/file/d/1G1V3LPTWxYFVXWd9fyyi7MiaDN0YeAJN/view?usp=sharing)):**

**Movie S1.**
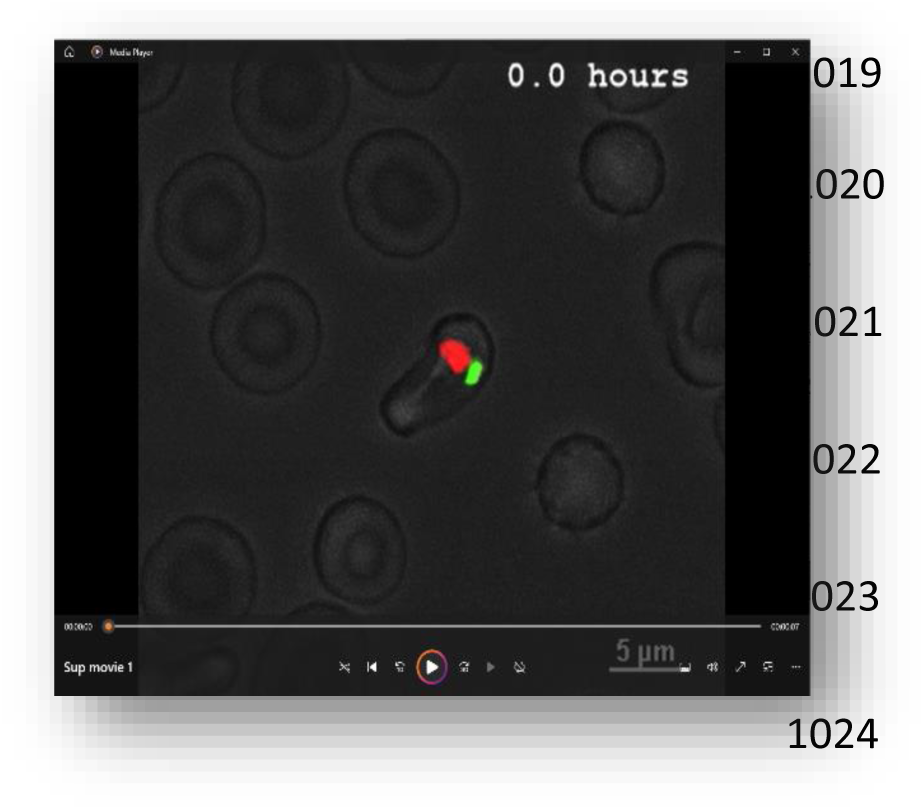
4D live imaging of *P. falciparum* reporter line, demonstrating nuclear divisions and apicoplast biogenesis. Representing single-cell crop MIP images acquired by live-cell fluorescence long-term 3D microscopy using H2B-mRuby-2TA-TP-mNeon line. Synchronized cells were prepared and imaged for 24 h at 15 min intervals, as described in Material and Methods using spinning disk microscopy. Raw data were processed to create single-cell crops (XYZ) at relevant time points (T1 to Tx (x=2 time points after egress)) for further measurements of volume, surface area and number of objects for each fluorescent channel.

**Sup. Movie S2 ([link](https://drive.google.com/file/d/1m1VEPQ9YilJw0L76NyKZUW94TljvR4Xx/view?usp=drive_link)):**

**Movie S2.**
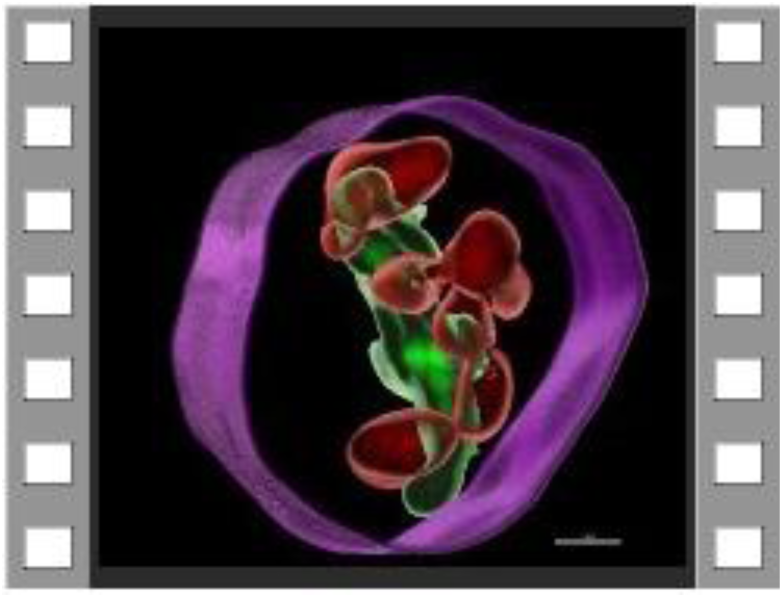
3D visualization of single-cell time-lapse crop images by Imaris. Automated surface masks were created for nucleus (red) and apicoplast (green) using fluorescent channels (561 and 488, accordingly) with appropriate thresholds. Cell membrane contour (purple) was manually drawn based on first time-point image acquired by DIC channel.

**Movie S3 ([link](https://drive.google.com/file/d/1MFMyzt9rWV5ih4gy--unNIJ25RVT9j2C/view?usp=drive_link))**

**Movies S3.**
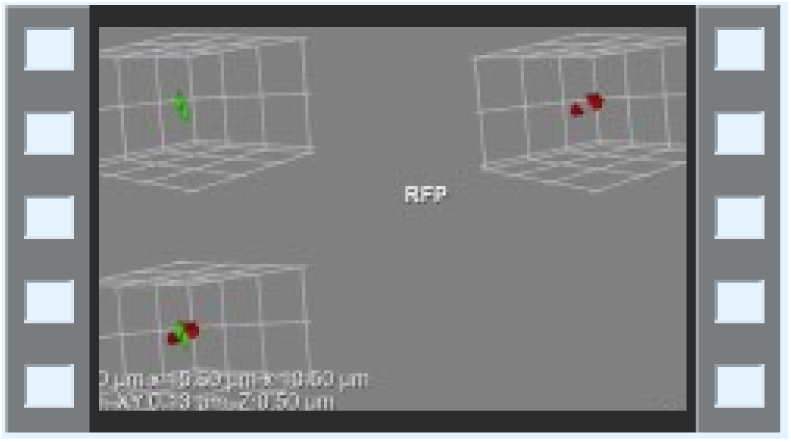
3D Live Apicoplast morphologies and association with nucleus. 3D volume rendering of a single time point show distinct apicoplast morphologies (green) and unique association with a dividing nucleus (red).

**Movie S4 ([link](https://drive.google.com/file/d/1WFEOO2LEwxxkArrCyBN4AHZzer1_M2dN/view?usp=drive_link))**

**Movies S4.**
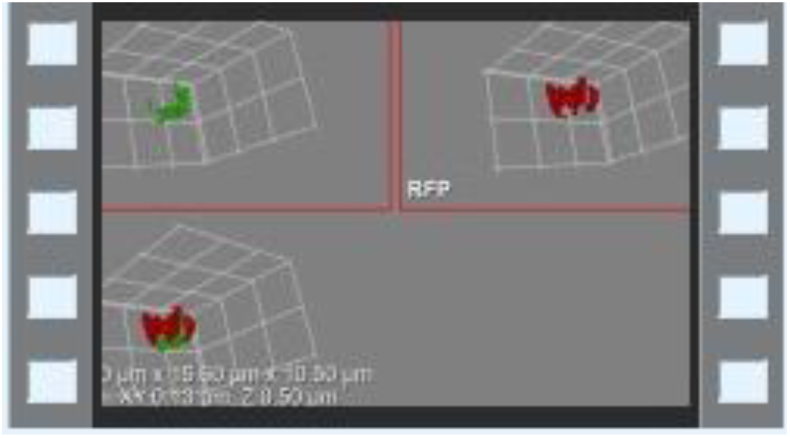
3D Live Apicoplast morphologies and association with nucleus. Branching apicoplast partly associated with multiple nuclei at later stages.

**Movie S5 ([link](https://drive.google.com/file/d/1RgDq6RZnxAPzLUo3ybAu0VT9xL40n3Ps/view?usp=drive_link))**

**Movies S5.**
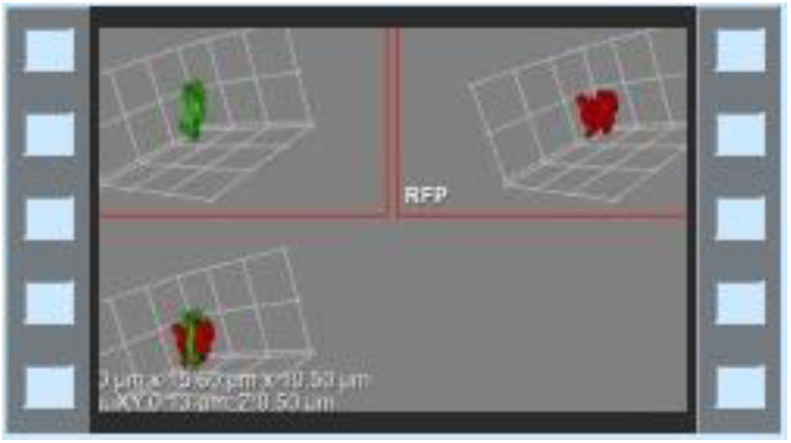
3D Live Apicoplast morphologies and association with nucleus. Crown morphology characterize an apicoplast stretched over multiple nuclei just before division.

**Movie S6 ([link](https://drive.google.com/file/d/1Hqjse_CSbFjvv0D4d6WqKvl_Nj4KqrT4/view?usp=drive_link))**

**Movies S6.**
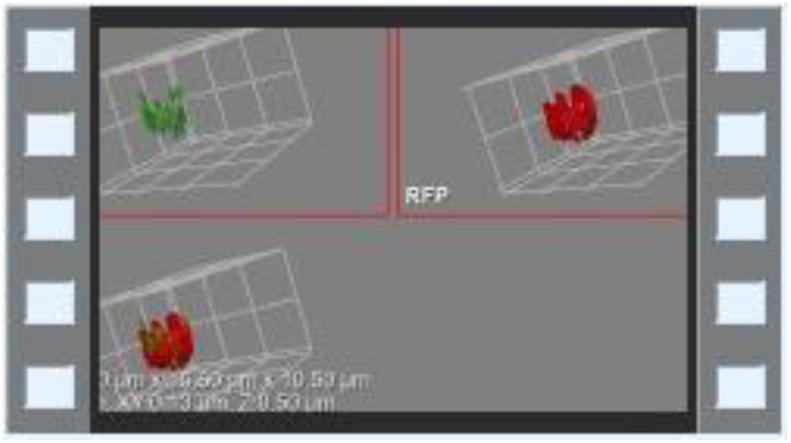
3D Live Apicoplast morphologies and association with nucleus. Apicoplast divides and closely associates with each new nucleus in the cell. Z-projections of 21 slices of 0.5μm. Brightness and contrast adjusted for better visibility.

**Sup. Movie S7 ([link](https://drive.google.com/file/d/1OIfcO09g5VYakMtsCkQ-CK8MTpqQpm0x/view?usp=drive_link))**

**Movie S7.**
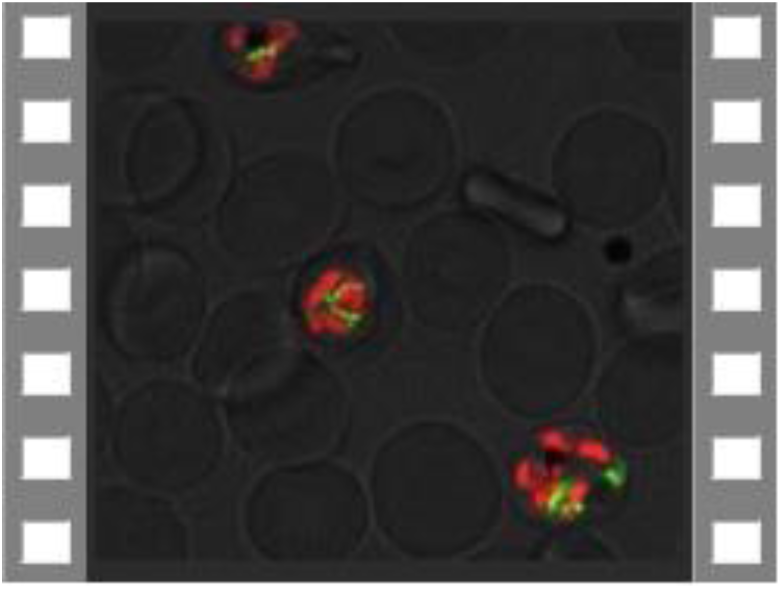
4D live imaging of *P. falciparum* H2B-mRuby-T2A-TP-mNeon reporter line. A large imaged field demonstrating co-development of nuclear divisions and apicoplast biogenesis. Long-term fluorescence live-cell microscopy show detailed apicoplast (green) and nucleus (red) morphologies and conformations throughout parasite’s life cycle in 3D (volume visualizations). Microscopy Technique: Spinning disk confocal microscopy. Images were captured at 100x as Z-stacks (total 21 slices of 0.5 μm each) with 15 min time intervals for 20 h. Scale bars, 2.5 μm.

**Sup. Movie S8 ([link](https://drive.google.com/file/d/1Fb8inrWaEJBCWAqgMitrCpxvb4K5tf8K/view?usp=drive_link))**

**Movie S8.**
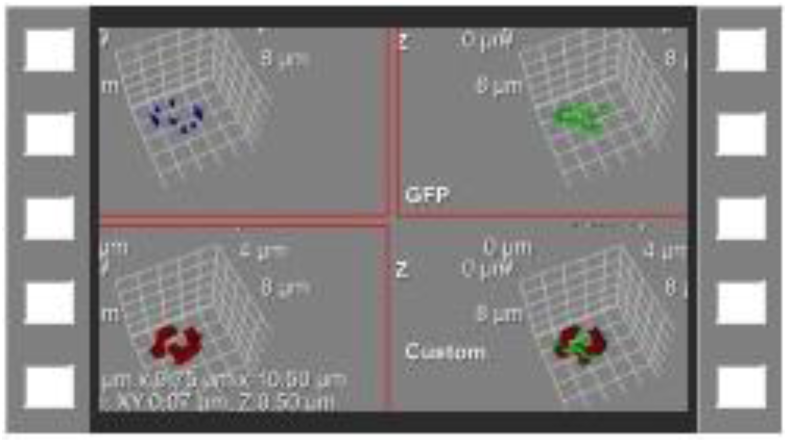
Tubulin mediates apicoplast-nucleus association. Live staining with Tubulin Tracker Deep Red along with live 4D microscopy reveals gradually growing association of apicoplast with nucleus-tubulin pairs at CP sites over time. 3D volume rendering of a time-lapse microscopy show apicoplast morphologies (green) and its association with a dividing nucleus (red) through binding to tubulin (blue) component of CP. Z-projections of 21 slices of 0.5μm. Brightness and contrast adjusted for better visibility.

**Sup. Movie S9 ([link](https://drive.google.com/file/d/1D-ZT8RV24oxSn5IKyqZ2lLEGp8XVN5e0/view?usp=drive_link))**

**Movie S9.**
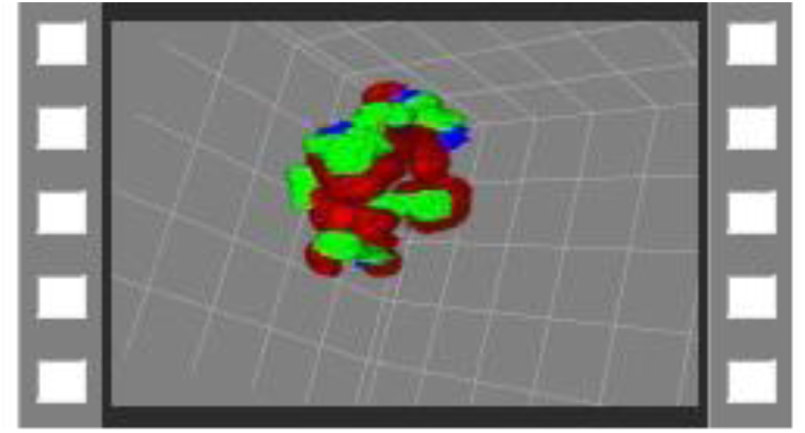
Apicoplast Crown morphology. 360^°^ rotation 3D volume image show a close association of a single large apicoplast (green) with multiple nuclei (red) by attachment to tubulin (blue) component of CP. Z-projections of 21 slices of 0.5μm. Brightness and contrast adjusted for better visibility.

**Sup. Movie S10 ([link](https://drive.google.com/file/d/1fVDvfpR-MA6znnKOrLRJNU9H6-Jv2CPU/view?usp=drive_link))**

**Movie S10.**
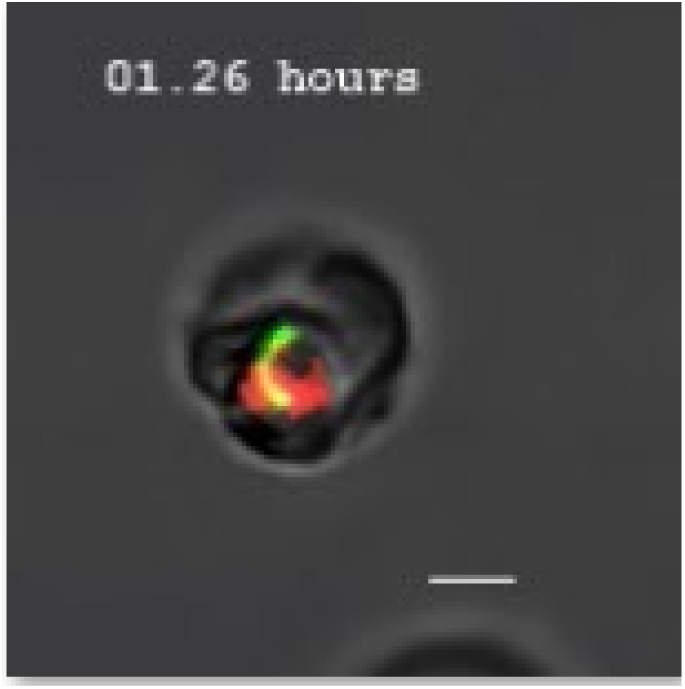
Aphidicolin arrest apicoplast biogenesis at early stage. Representing single-cell crop MIP images acquired by live 4D microscopy using synchronized H2B-mRuby-2TA-TP-mNeon line treated with 2μM Aphidicolin. Nuclear divisions inhibited at 1-2n state as well as apicoplast biogenesis failed to progress beyond elongation. Finally, the apicoplast stuck with an atypical morphology. Scale bar is 2.5μM.

**Sup. Movie S11 ([link](https://drive.google.com/file/d/1Go3SuLa4JeOyp1lSgN7X7O93rey192XO/view?usp=drive_link))**

**Movie S11.**
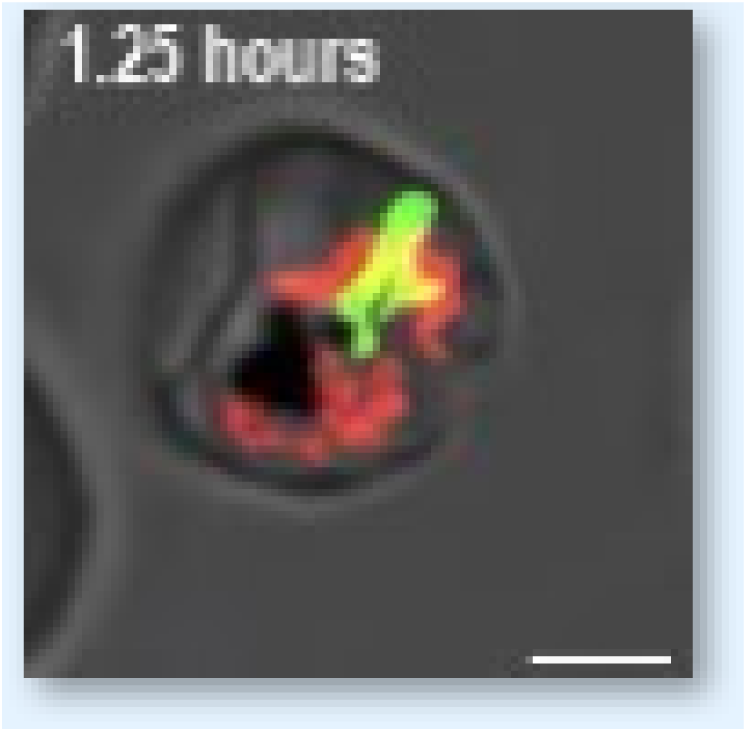
CIP affect apicoplast biogenesis at crown stage. Representing single-cell crop MIP images acquired by live 4D microscopy using synchronized H2B-mRuby-2TA-TP-mNeon line treated with 8μM CIP+IPP. Nuclear divisions proceed normally, while apicoplast failed to encircle all nuclei, resulting in impaired apicoplast division and egress. Scale bar is 2.5μM.

