## Supplementary figures and images for "Organelle Development and Inheritance are Driven by Independent Nuclear and Organellar Mechanisms in Malaria Parasites"

### Sup Fig 1

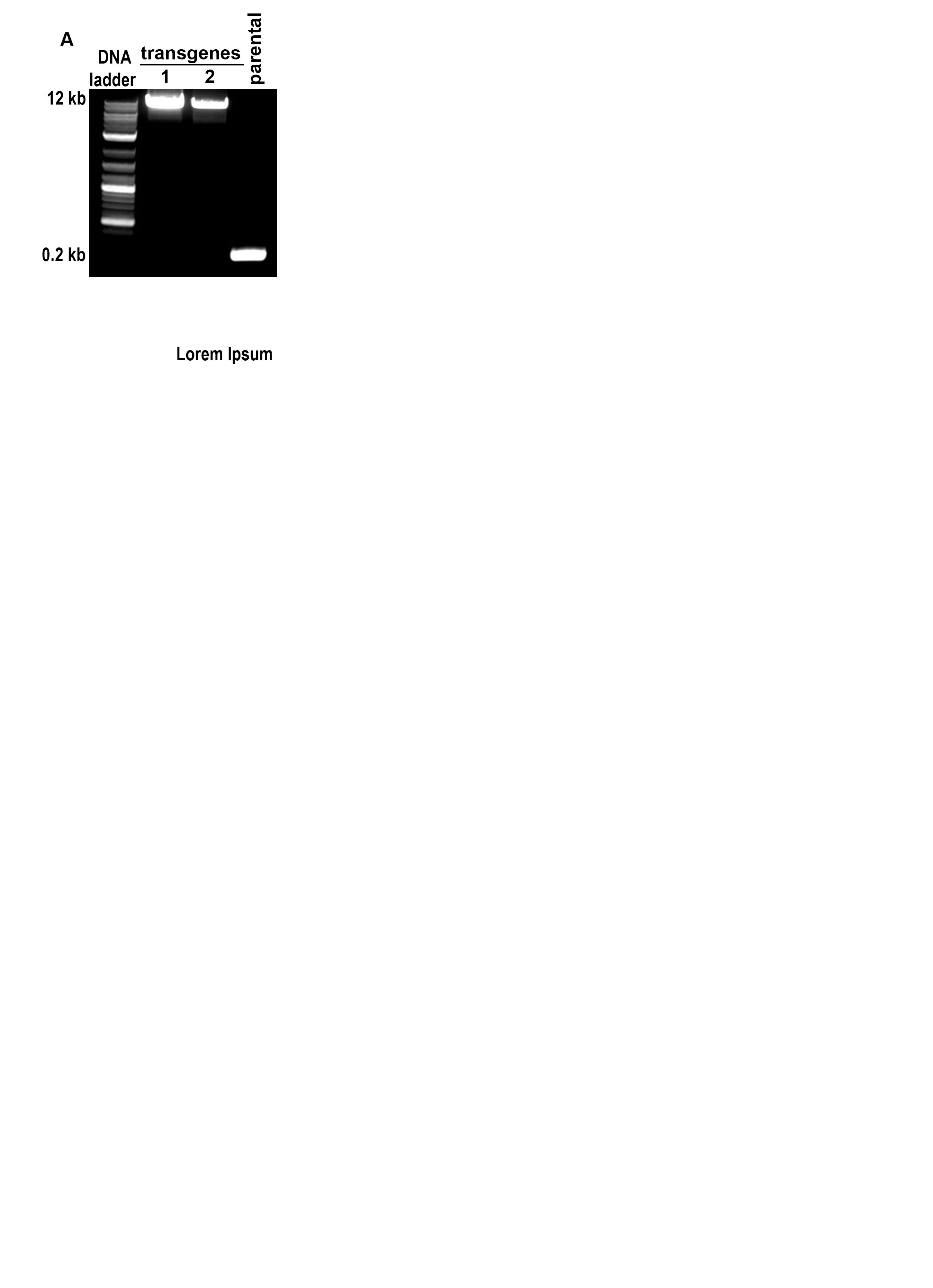

### Sup Fig 2

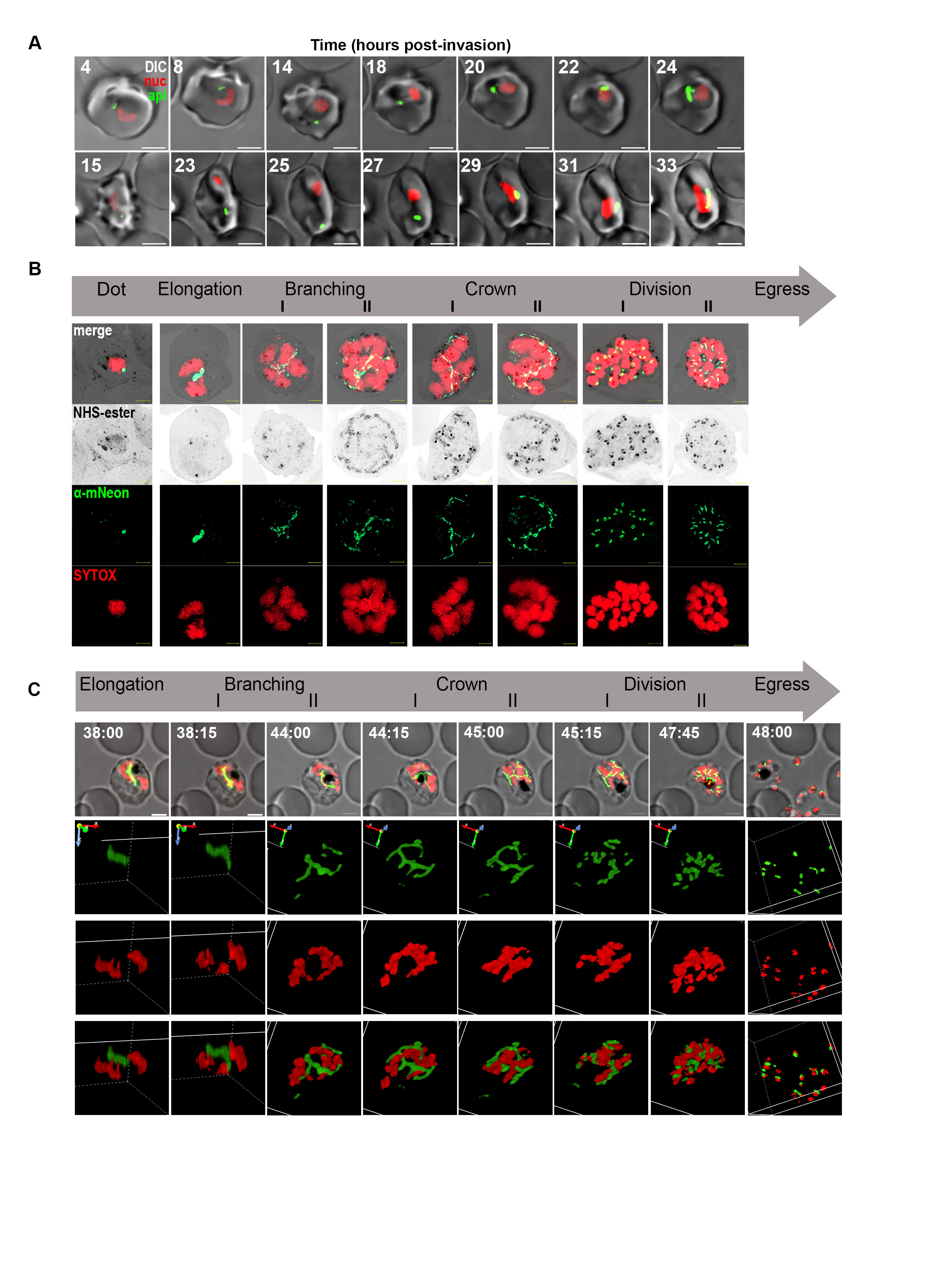

### Sup Fig 3

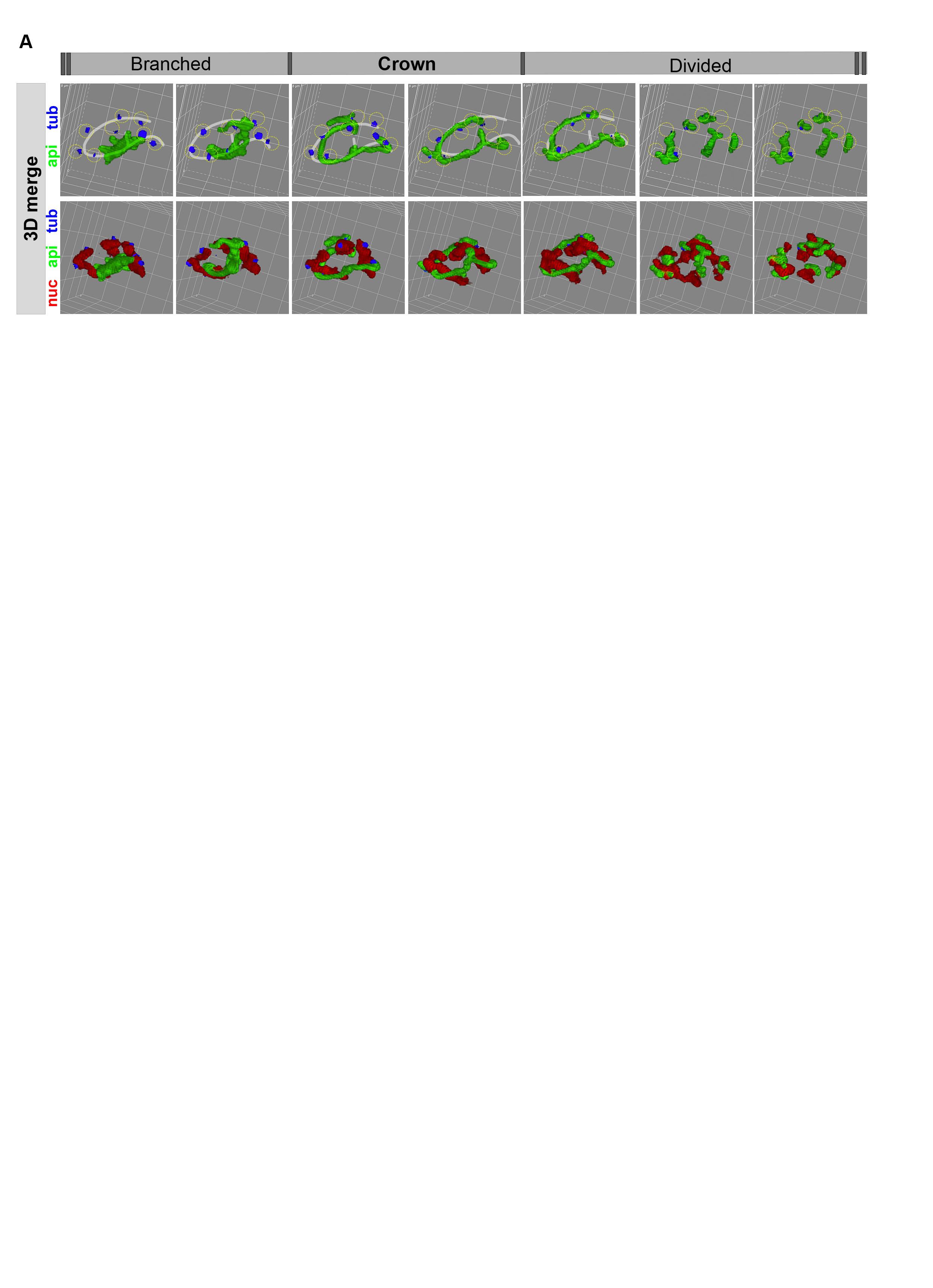

### Sup Fig 4

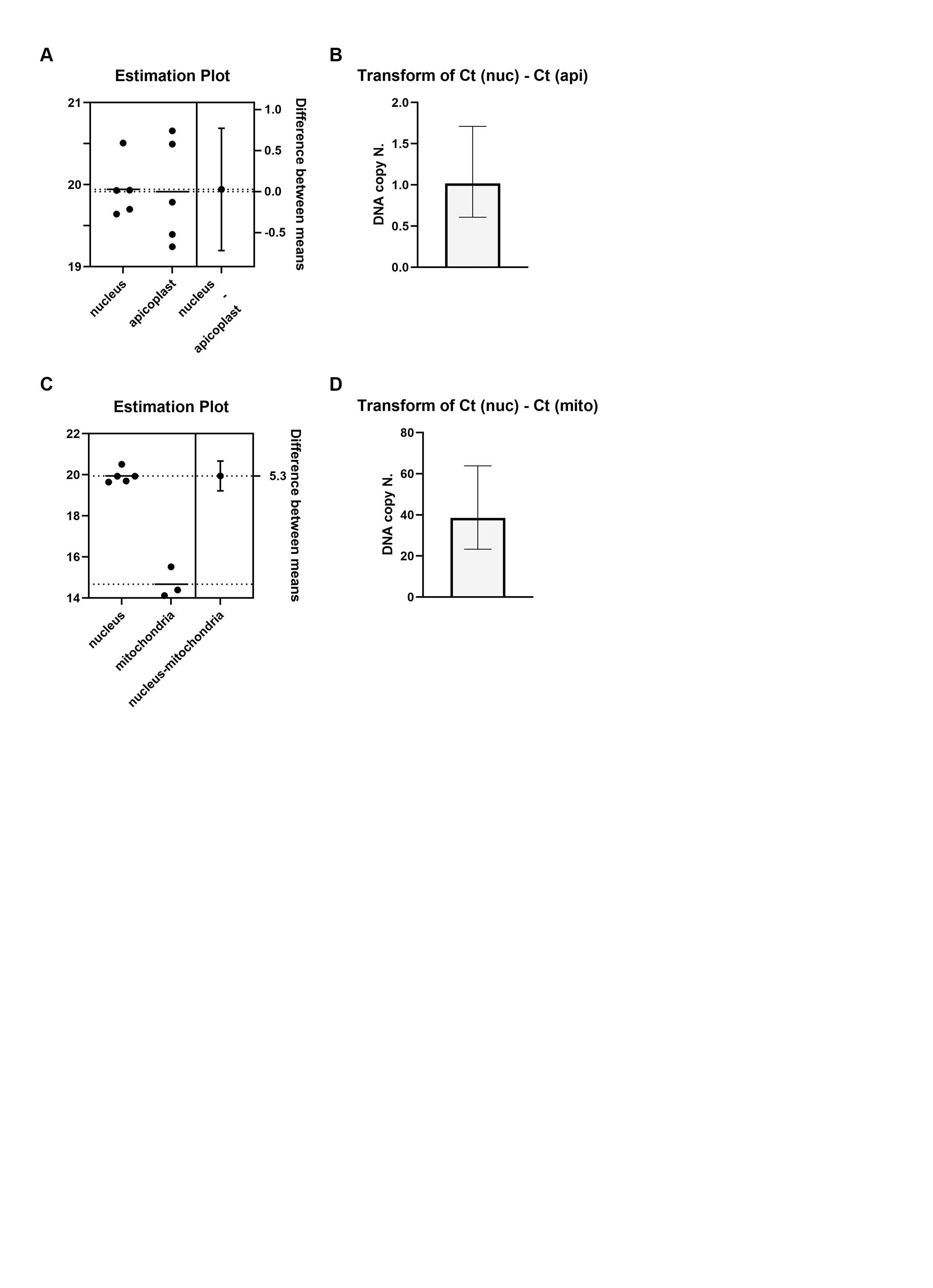

### Sup Fig 5

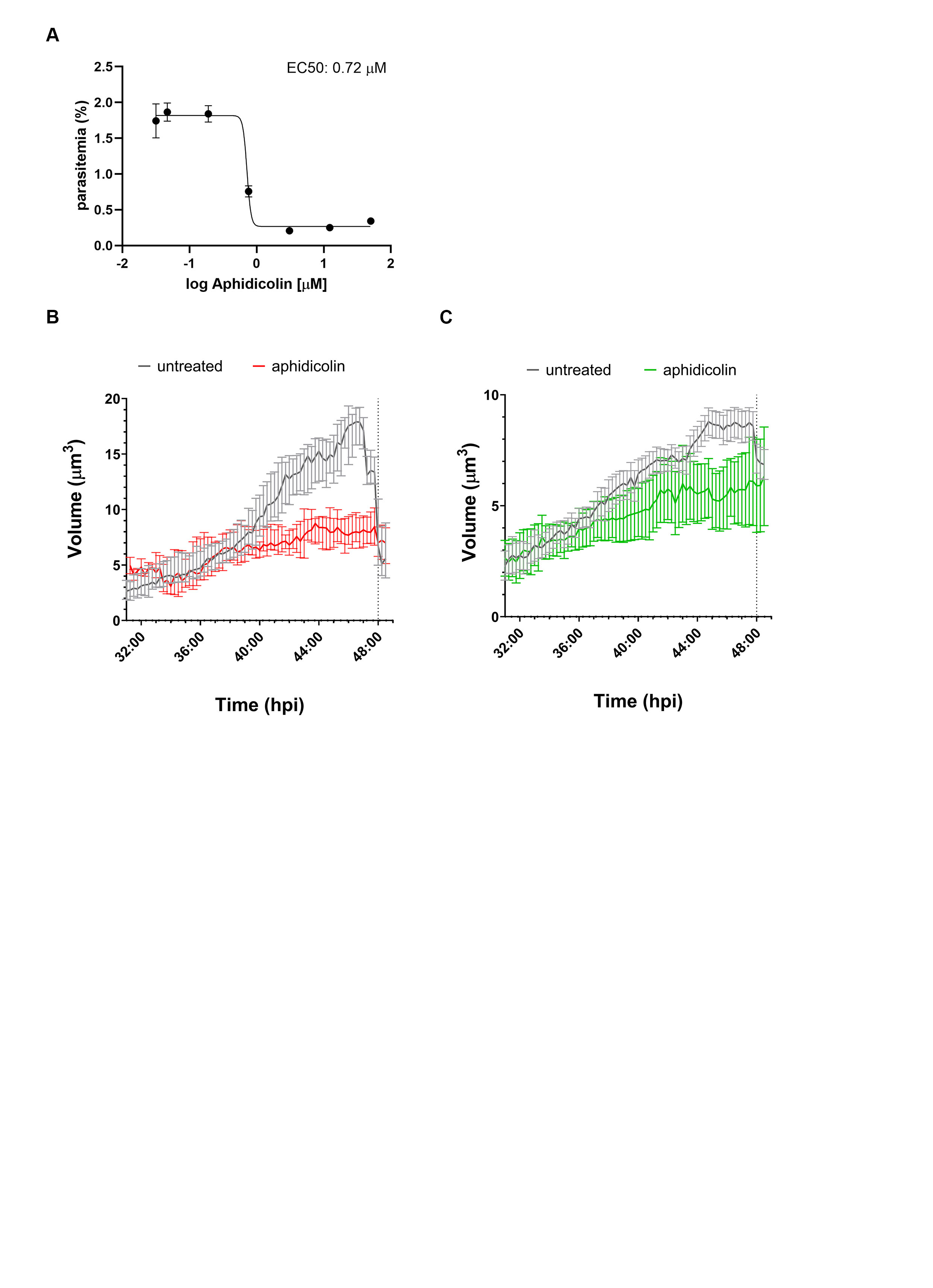

### Sup Fig 6

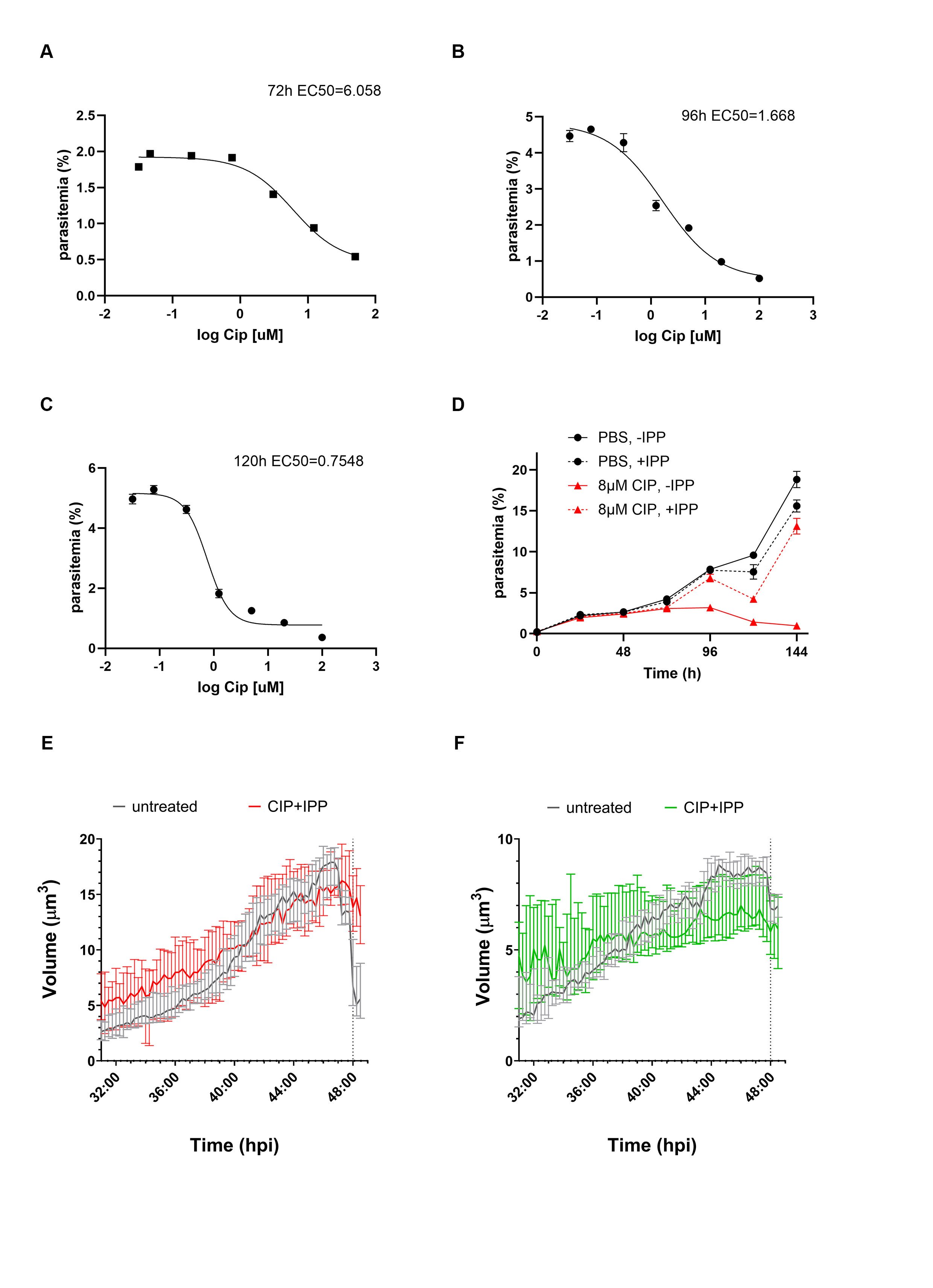
