## Supplementary material for "Organelle Development and Inheritance are Driven by Independent Nuclear and Organellar Mechanisms in Malaria Parasites": Sup Movies links

**Sup. Movie S1 (link):**

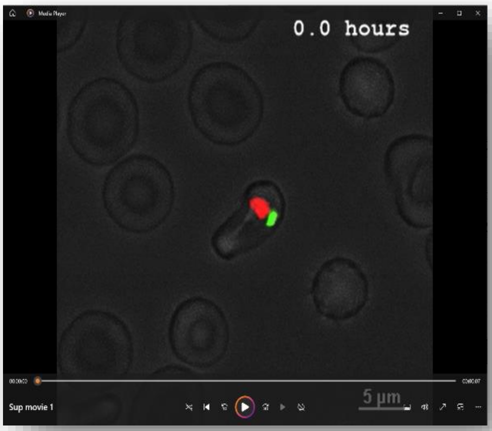


**Movie S1. 4D live imaging of *P. falciparum* reporter line, demonstrating nuclear divisions and apicoplast biogenesis**. Representing single-cell crop MIP images acquired by live-cell fluorescence long-term 3D microscopy using H2B-mRuby-2TA-TP-mNeon line. Synchronized cells were prepared and imaged for 24 h at 15 min intervals, as described in Material and Methods using spinning disk microscopy. Raw data were processed to create single-cell crops (XYZ) at relevant time points (T1 to Tx (x=2 time points after egress)) for further measurements of volume, surface area and number of objects for each fluorescent channel.

**Sup. Movie S2 (link):**

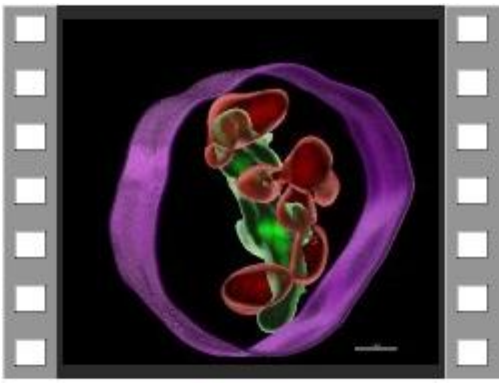

**Movie S2. 3D visualization of single-cell time-lapse crop images by Imaris.** Automated surface masks were created for nucleus (red) and apicoplast (green) using fluorescent channels (561 and 488, accordingly) with appropriate thresholds. Cell membrane contour (purple) was manually drawn based on first time-point image acquired by DIC channel.

**Sup. Movie S3-6**

**Movie S3 (link)**

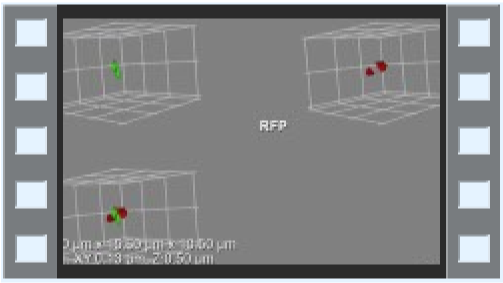


**Movie S4 (link)**

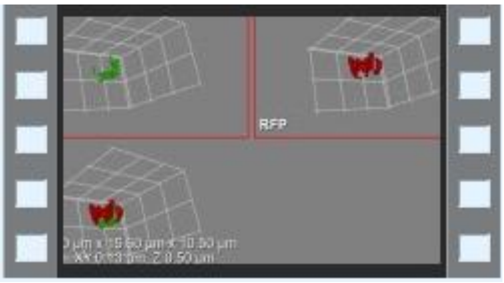


**Movie S5 (link)**

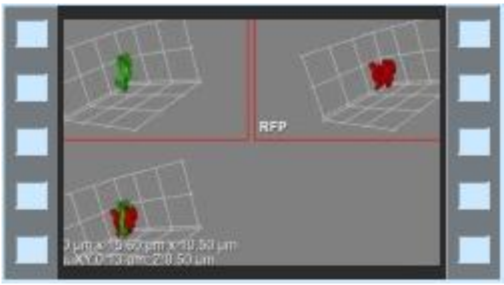


**Movie S6 (link)**

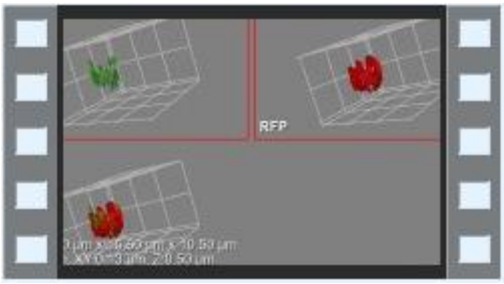


**Movies S3-6**. **3D Live Apicoplast morphologies and association with nucleus**. 3D volume rendering of a single time point show distinct apicoplast morphologies (green) and unique association with a dividing nucleus (red). **S3**. Elongated apicoplast at the onset of biogenesis is not associated with newly divided nuclei (2-3n). **S4**. Branching apicoplast partly associated with multiple nuclei at later stages. **S5**. Crown morphology characterize an apicoplast stretched over multiple nuclei just before division. **S6**. Apicoplast divides and closely associates with each new nucleus in the cell. Z-projections of 21 slices of 0.5μm. Brightness and contrast adjusted for better visibility.

**Sup. Movie S7 (link)**

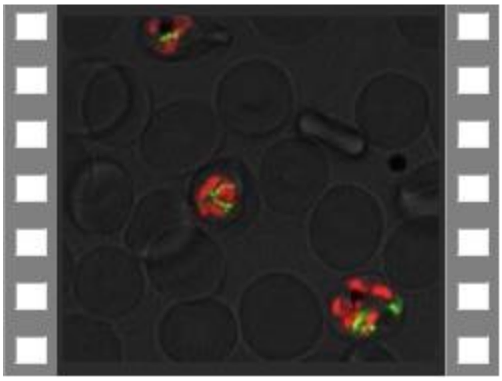

**Movie S7. 4D live imaging of *P. falciparum* H2B-mRuby-T2A-TP-mNeon reporter line.** A large imaged field demonstrating co-development of nuclear divisions and apicoplast biogenesis. Long-term fluorescence live-cell microscopy show detailed apicoplast (green) and nucleus (red) morphologies and conformations throughout parasite's life cycle in 3D (volume visualizations). Microscopy Technique: Spinning disk confocal microscopy. Images were captured at 100x as Z-stacks (total 21 slices of 0.5 µm each) with 15 min time intervals for 20 h. Scale bars, 2.5 µm.

**Sup. Movie S8 (link)**

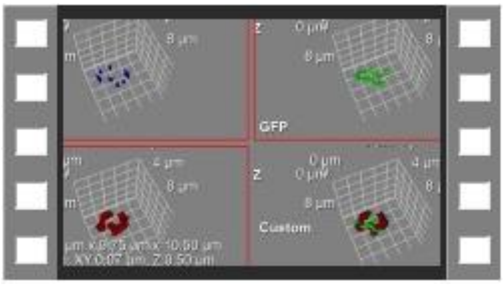


**Movie S8. Tubulin mediates apicoplast-nucleus association.** Live staining with Tubulin Tracker Deep Red along with live 4D microscopy reveals gradually growing association of apicoplast with nucleus-tubulin pairs at CP sites over time. 3D volume rendering of a time-lapse microscopy show apicoplast morphologies (green) and its association with a dividing nucleus (red) through binding to tubulin (blue) component of CP. Z-projections of 21 slices of 0.5µm. Brightness and contrast adjusted for better visibility.

**Sup. Movie S9 (link)**

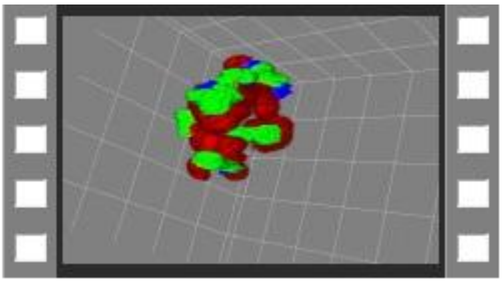

**Movie S9. Apicoplast Crown morphology.** $360^0$ rotation 3D volume image show a close association of a single large apicoplast (green) with multiple nuclei (red) by attachment to tubulin (blue) component of CP. Z-projections of 21 slices of 0.5μm. Brightness and contrast adjusted for better visibility.

**Sup. Movie S10 (link)**

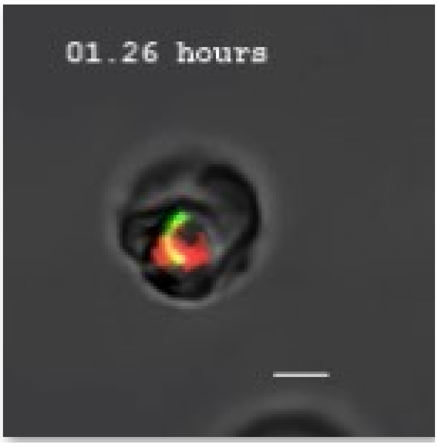


**Movie S10. Aphidicolin arrest apicoplast biogenesis at early stage.** Representing single-cell crop MIP images acquired by live 4D microscopy using synchronized H2B-mRuby-2TA-TP-mNeon line treated with 2μM Aphidicolin. Nuclear divisions inhibited at 1-2n state as well as apicoplast biogenesis failed to progress beyond elongation. Finally, the apicoplast stuck with an atypical morphology. Scale bar is 2.5μM.

**Sup. Movie S11 (link)**

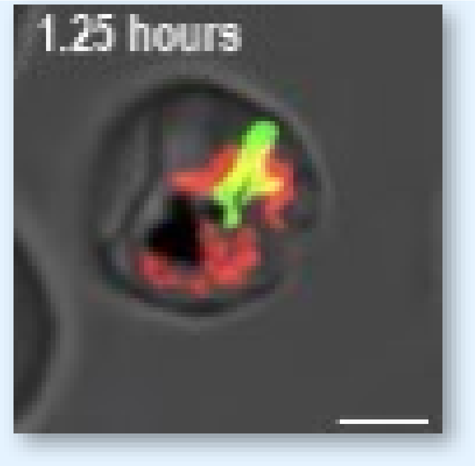


**Movie S11. CIP affect apicoplast biogenesis at crown stage.** Representing single-cell crop MIP images acquired by live 4D microscopy using synchronized H2B-mRuby-2TA-TP-mNeon line treated with 8μM CIP+IPP. Nuclear divisions proceed normally, while apicoplast failed to encircle all nuclei, resulting in impaired apicoplast division and egress. Scale bar is 2.5μM.
